# A combined transcriptional and dynamic roadmap of single human pancreatic endocrine progenitors reveals proliferative capacity and differentiation continuum

**DOI:** 10.1101/2021.12.15.472220

**Authors:** Belin Selcen Beydag-Tasöz, Joyson Verner D’Costa, Lena Hersemann, Federica Luppino, Yung Hae Kim, Christoph Zechner, Anne Grapin-Botton

## Abstract

Basic helix-loop-helix genes, particularly proneural genes, are well-described triggers of cell differentiation, yet limited information exists on their dynamics, notably in human development. Here, we focus on Neurogenin 3 (*NEUROG3*), which is crucial for pancreatic endocrine lineage initiation. Using a double reporter to monitor endogenous *NEUROG3* transcription and protein expression in single cells in 2D and 3D models of human pancreas development, we show peaks of expression for the RNA and protein at 22 and 11 hours respectively, approximately two-fold slower than in mice, and remarkable heterogeneity in peak expression levels all triggering differentiation. We also reveal that some human endocrine progenitors proliferate once, mainly at the onset of differentiation, rather than forming a subpopulation with sustained proliferation. Using reporter index-sorted single-cell RNA-seq data, we statistically map transcriptome to dynamic behaviors of cells in live imaging and uncover transcriptional states associated with variations in motility as NEUROG3 levels change, a method applicable to other contexts.

## Introduction

Endocrine pancreas development has been under scrutiny in the last decades, and understanding its formation has enabled the engineering of β-cells from human pluripotent stem cells (hPSCs) for cell therapy of diabetes mellitus, a complex metabolic disease affecting more than 450 million people worldwide (International Diabetes Federation, 2019). Improvements over the control of differentiation and functionality of β-cells depend on a better understanding of endocrinogenesis, starting with the common endocrine progenitors that can give rise to all five pancreatic endocrine subtypes, namely insulin-producing β-cells, glucagon-producing α-cells, somatostatin-producing δ-cells, ghrelin-producing ε-cells, and pancreatic polypeptide-producing PP-cells.

Pancreatic organogenesis and lineage hierarchy are generally considered to be similar between humans and mice (Jennings et al., 2013; Petersen et al., 2018). In the context of the endocrine differentiation, transient expression of the basic helix-loop-helix (bHLH) transcription factor Neurogenin 3 (*NEUROG3*) is the trigger of pancreatic endocrine lineage formation in both species (Gradwohl et al., 2000; Gu et al., 2002; Heller et al., 2005; Johansson et al., 2007; McGrath et al., 2015; Ohsie et al., 2009; Pinney et al., 2011; Rubio-Cabezas et al., 2011; Schwitzgebel et al., 2000). *Neurog3* knockout mice fail to produce any pancreatic endocrine cells and die postnatally from diabetes mellitus (Gradwohl et al., 2000), and in almost all described cases of severe *NEUROG3* mutations in humans, patients developed diabetes mellitus early in life (Pinney et al., 2011; Rubio-Cabezas et al., 2011; Solorzano-Vargas et al., 2020). In contrast, some other aspects of endocrine pancreas development are not conserved between humans and mice. One example is the biphasic *Neurog3* expression observed in mice (Villasenor et al., 2008), which does not seem to exist in humans (Jennings et al., 2013; Salisbury et al., 2014). Therefore, studying human pancreatic development is essential to uncover interspecies differences. We previously showed that *Neurog3* reaches its peak expression level in about 12 hours in single cells in mouse embryonic pancreas explants (Kim et al., 2015). In comparison, the duration of *NEUROG3* transcript and protein expression in single cells has not been previously measured in a human system.

With the increased levels of NEUROG3, the mouse endocrine progenitors start to exit the cell cycle, as NEUROG3 is known to induce the expression of cell-cycle inhibitors such as cyclin dependent kinase inhibitor 1a (*Cdkn1a*) and p21 (RAC1) activated kinase 3 (*Pak3*) (Miyatsuka et al., 2011; Piccand et al., 2014). However, recent studies have shown that cells with low levels of *Neurog3* mRNA can themselves proliferate, but the extent of this proliferation is not clear (Bechard et al., 2016). Whether such proliferative low *NEUROG3-* expressing cells exist in humans as well has not yet been addressed, but represents a key question as a proliferative population committed to the endocrine lineage would be interesting to propagate with the aim of producing β-cells for cell therapy.

In this study, we set out to determine the kinetics of human endocrine progenitor differentiation and investigate their proliferation capacity using 2-dimensional (2D) and 3-dimensional (3D) *in vitro* human pancreatic differentiation and culturing methods to model early steps of human pancreatic endocrine development (Petersen et al., 2017; Rezania et al., 2014). Moreover, we engineered a tagged human NEUROG3 protein that enabled us to determine the duration of NEUROG3 expression in human endocrine progenitors. Our construct also monitors *Neurog3* transcription, which can be compared to similar reporters in mice (Kim et al., 2015). Mapping live imaging to the deep single-cell RNA sequencing at the emergence point of endocrine progenitors allowed us to correlate cellular dynamics to transcriptomics and uncovered differences in *NEUROG3*+ cell behavior and movement speed. This work highlights the importance of using human systems to study pancreatic endocrine cell differentiation to better understand these cells and human pancreas development, critical for *in vitro* endocrine cell production for diabetes mellitus therapy.

## Results

### Generation of a dual reporter line monitoring *NEUROG3* transcription and protein in a human system in real-time

To visualize the first steps of endocrine pancreas specification in a human system, we constructed a dual reporter human embryonic stem cell (hESC) line using CRISPR-Cas9 gene editing. This allows monitoring *NEUROG3* transcription and protein using one of the alleles at the endogenous *NEUROG3* locus (Figure 1A). This reporter line produces a fusion protein of NEUROG3 with TagRFP-T (RFP) to live-monitor the human NEUROG3 protein and EGFP (GFP) targeted to the nucleus to track the transcription from the *NEUROG3* locus simultaneously. This strategy makes it possible to continue to follow the cells that turned off *NEUROG3* expression due to the slow degradation of GFP (Petersen et al., 2017). Recent studies indicate that human NEUROG3 protein can be degraded relatively quickly (Krentz et al., 2017).

**Figure 1:**
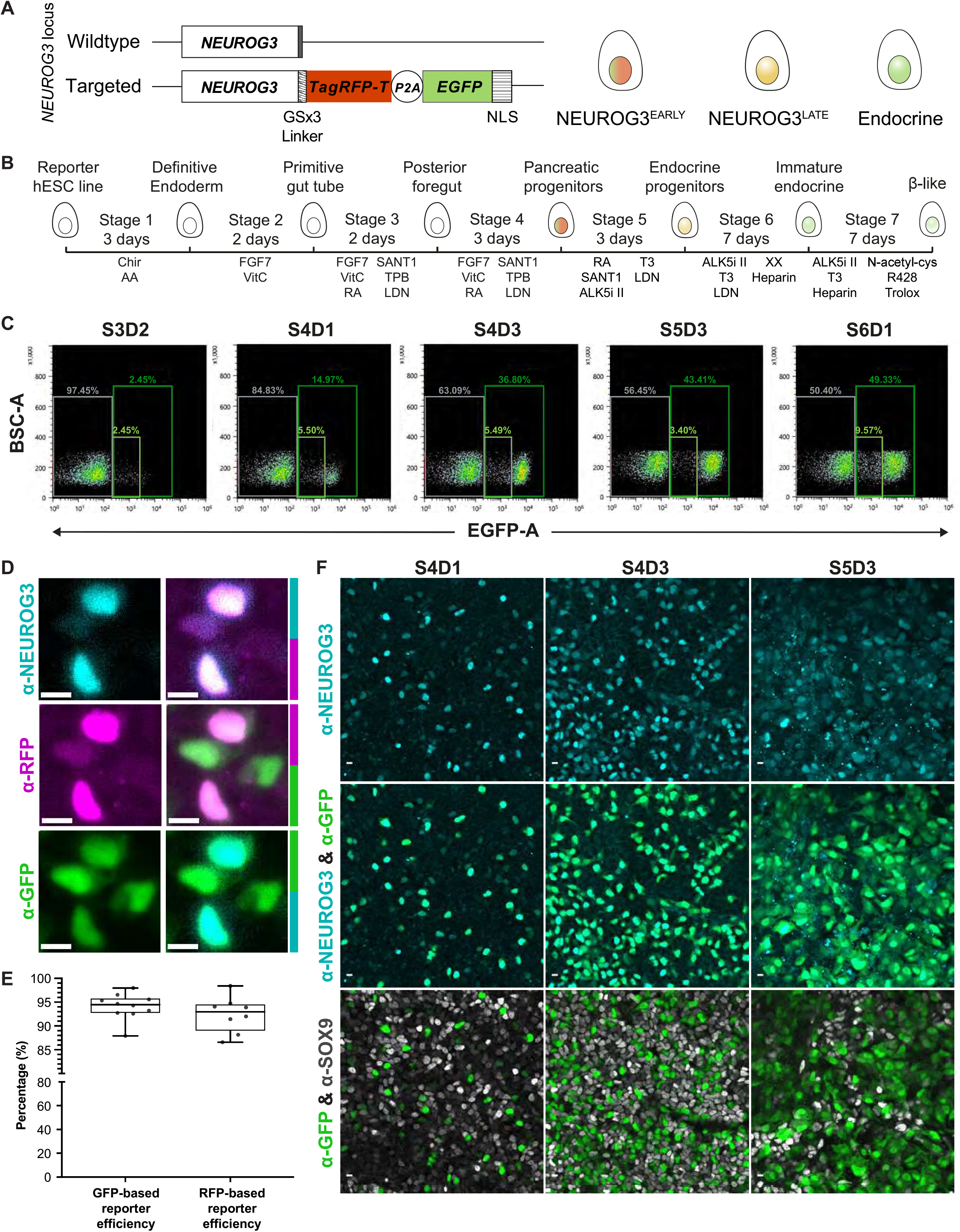
Construction and characterization of a reliable dual hESC-based reporter line monitoring human *NEUROG3* transcription and protein expression. **1A:** Schematic representation of the dual fluorescent reporter construct introduced in the endogenous *NEUROG3* locus and the fluorescence progression of *NEUROG3*-expressing cells. The upper lane shows the wildtype *NEUROG3* allele, and the lower lane shows the targeted *NEUROG3* locus where the reporter sequence encoding *TagRFP-T* with a flexible GSx3 linker, a self-cleaving peptide (*P2A*), and *EGFP* tagged with a nuclear localization signal (NLS) was introduced before the stop codon of the *NEUROG3* coding sequence. With this setup, *NEUROG3*-expressing cells are nuclear-labeled with red and green fluorescent proteins. As *NEUROG3* is transcribed, EGFP accumulates in the cell, and after *NEUROG3* is downregulated, longer-living EGFP allows tracing of these cells using the green fluorescence. **1B:** Illustration of the *in vitro* pancreatic differentiation protocol starting from hESCs. The individual stages are indicated along with the time scale and added components. **1C:** Progression of the GFP expression from Stage 3 Day 2 to Stage 6 Day 1 of the *in vitro* pancreatic differentiation analyzed by flow cytometry. Back scatter (BSC) versus GFP profiles obtained from cells collected at different stages of the *in vitro* pancreatic differentiation protocol are shown. Light green gate marks GFP-low cells, while dark green gate marks all GFP+ cells. Grey gate indicates GFP-negative cells. **1D:** Representative immunofluorescence microscopy images showing NEUROG3, GFP, and RFP expression in differentiated human pancreatic cells. Anti-NEUROG3 (cyan), anti-RFP (magenta), and anti-EGFP (green). Left panels show single colors, while the right panels show co-expression of pairs. Scale bars, 10 μm. **1E:** Boxplots showing the efficiency of the dual *NEUROG3* reporter line calculated by image-based quantification of the percentage of co-expression between NEUROG3 and GFP or RFP within the anti-NEUROG3-positive compartment. *N* = 3 and *n* = 10 for GFP, and *N* = 1 and *n* = 8 for RFP. **1F:** Representative immunofluorescence microscopy images showing NEUROG3 and SOX9. Anti-NEUROG3 (cyan), anti-GFP (green), and anti-SOX9 (gray). Scale bars, 10 μm. S: Stage, D: Day

Using a 7-stage *in vitro* human pancreatic endocrine differentiation protocol, we differentiated our hESC dual *NEUROG3* reporter line into the pancreatic lineage (Figure 1B). The karyotype analyses showed no abnormality (Figure S1A). The GFP and RFP-positive endocrine progenitors started to emerge during Stage 4 of the differentiation protocol, with their percentage increasing throughout Stages 4 and 5 (Figure 1C and S1B). Notably, the RFP signal was not detectable through flow cytometry, although it was detected by microscopy (Figure 1D). The reliability of our reporter was assessed by quantifying the percentage of cells expressing NEUROG3 protein that were also GFP or RFP-positive. We found that 92% and 94% of NEUROG3-positive cells also expressed RFP and GFP, respectively (Figures 1D-F). The reporters were also specific for NEUROG3-expressing cells as RFP was not detected in NEUROG3-negative cells (over 96% of RFP-positive cells also expressed NEUROG3). Our reporter line further showed comparable endocrine differentiation efficiency to the mother H1 hESC line (Figure S1C). Therefore, after validating our reporter line and differentiation platform, we set up our live imaging system to visualize newly emerging endocrine progenitor cells.

### *NEUROG3* expression levels are heterogeneous among human pancreatic cells

We started our live imaging experiment from the beginning of Stage 4 (Stage 3 Day 2), at the onset of *NEUROG3 expression* (Figures 1C-E), and performed the imaging over 5 days (Figure 2A). As our focus was on the initiation of *NEUROG3* expression, we traced the newly emerging *NEUROG3*+ cells. We observed a high degree of heterogeneity in the *NEUROG3* peak expression level and dynamics in human cells (Figures 2B and S2A-B). Human NEUROG3 fusion protein peaked around 11 hours after induction but did not go back to base levels within 26 hours of imaging (Figure 2C), so the duration of expression was addressed in the later experiments with a longer tracing span. Using the GFP transcriptional reporter, we compared the transcription dynamics from the human *NEUROG3* locus in individual cells to the published work in which mouse *Neurog3* promoter was used to drive turbo RFP expression (Kim et al., 2015). In human cells, *NEUROG3* transcription reached its peak in around 22 hours on average, whereas in the mouse cells, it reached it in 12 hours, highlighting a faster process in mice than in the human systems (Figure 2D). This observation indicated that human *NEUROG3* expression in each cell is slower and lasts longer than in the mouse, as shown with a less steep slope of transcription reporter signal accumulation. From the current imaging datasets, we moved on to investigating the behavior of endocrine progenitors.

**Figure 2:**
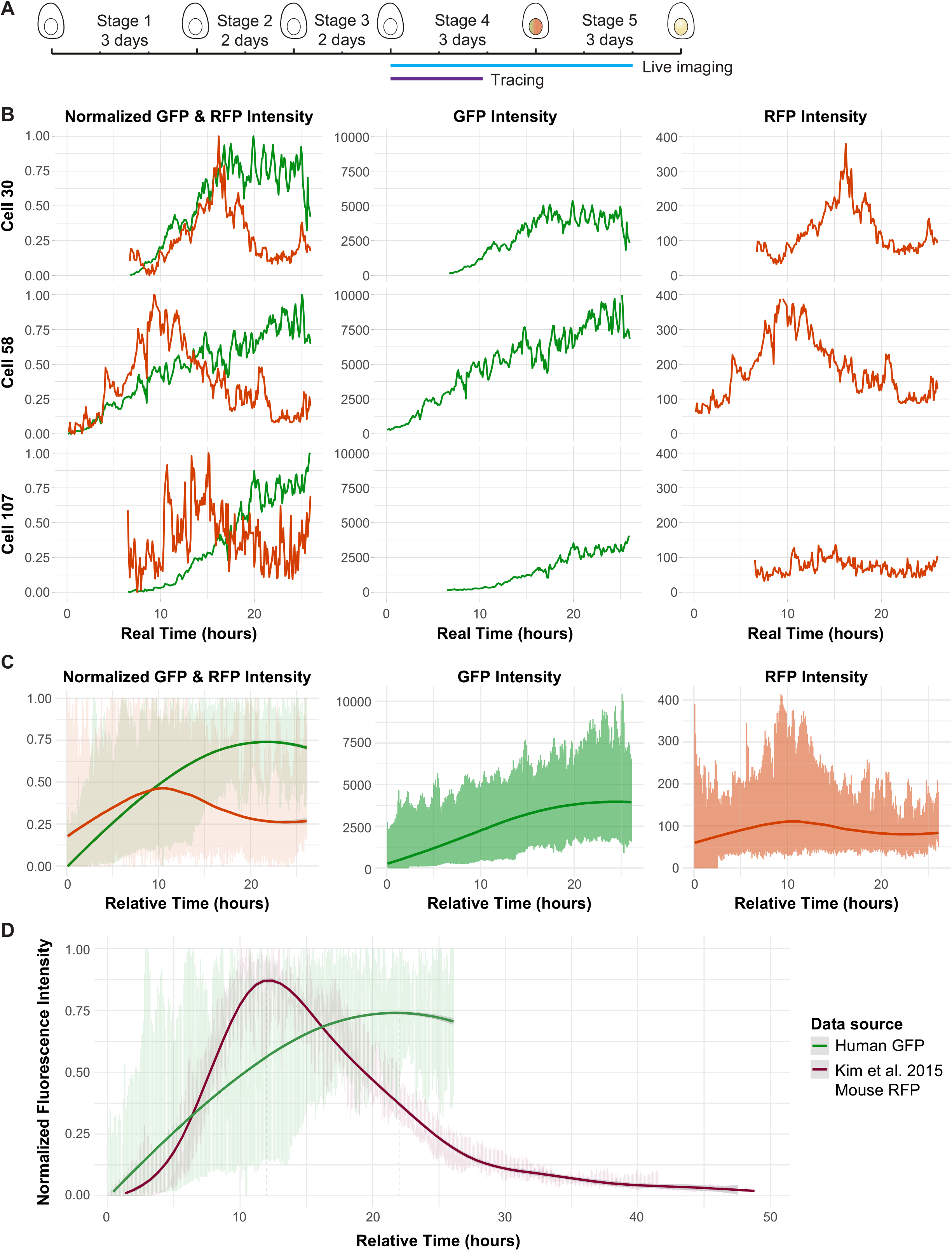
Live imaging of *NEUROG3*+ human pancreatic cells reveals heterogeneity of *NEUROG3* expression and a slower accumulation of transcriptional reporter compared to that of the mouse. **2A:** Schematic representation of the timing of live imaging and single-cell tracing over the *in vitro* pancreatic differentiation timeline. **2B:** Representative fluorescence tracing plots for three selected *NEUROG3*+ cells, showing GFP and RFP fluorescence intensities over time. On the left panel, fluorescence intensities were scaled between 0 and 1. **2C:** Plots showing fluorescent traces of all traced *NEUROG3*+ cells together. On the left panel, fluorescence intensities were scaled between 0 and 1. Dark green (GFP) and dark red (RFP) lines show smoothened mean intensities. The distribution represents individual values for each track per time point and the gray area shows the confidence interval. *N* = 3, *n* = 52 **2D:** Comparison of the dynamics of human *NEUROG3* transcription with the mouse *Neurog3* transcription from Kim et al., 2015. The green line shows smoothened mean GFP fluorescence intensities for the dual *NEUROG3* reporter, and the maroon line shows average mean RFP intensities for the mouse transcriptional *Neurog3* reporter. The distribution represents individual values for each track per time point and the gray area shows the confidence interval. *N* = 3, *n* = 52 for human GFP, and *n* = 6 for mouse RFP.

### Human endocrine progenitors can divide after *NEUROG3* onset

Since studies in mice have suggested that endocrine progenitors can proliferate (Bechard et al., 2016), we next addressed whether human endocrine progenitors could proliferate. Quantification from live imaging revealed that, on average, 4.8% of GFP+ cells divide in 26 hours, mainly at the onset of differentiation (Figures 3A and 3B). We did not notice a pattern in the timing (Figure 3C) or the location (Figure S3A) of the division events that might indicate a hotspot for a time or place. Even when we normalized the timing of the division to the onset of GFP, we did not find any correlation of the division time among dividing cells, but they were rather asynchronous (Figure 3C). In general, we noted that these proliferative endocrine progenitor cells tend to divide within 12 hours after *NEUROG3* induction marked by low GFP levels, with a few exceptions (Figures 3C and 3D).

**Figure 3:**
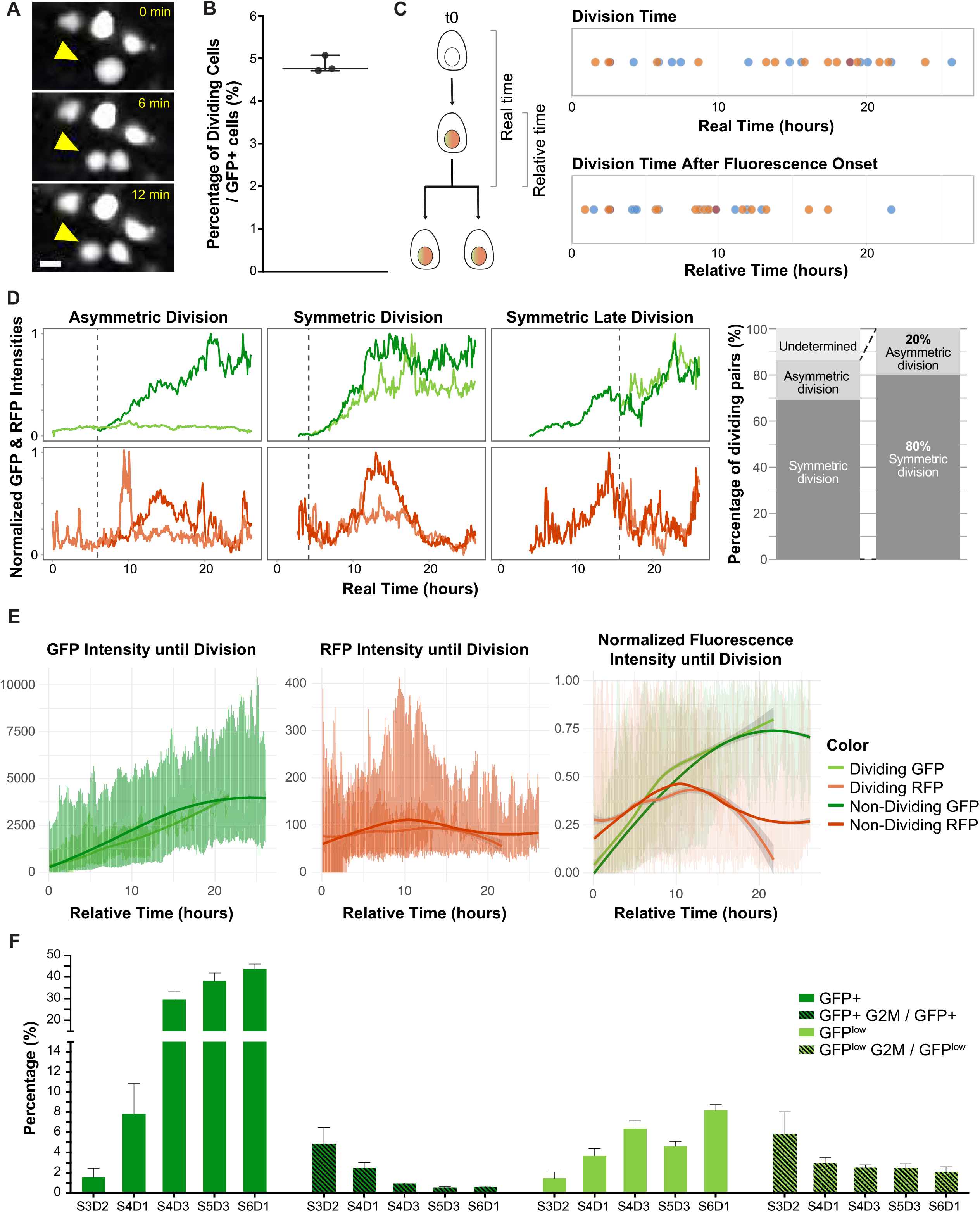
Single-cell profiling reveals that a subset of human *NEUROG3*+ progenitors are in the cell cycle. **3A:** Representative snapshot images showing division (arrowhead) in a GFP+ human pancreatic cell at 0, 6, and 12 minutes (relative time). Scale bars, 10 μm. **3B:** Boxplot showing the percentage of dividing cells within the GFP+ compartment from the live imaging experiments. Whiskers indicate the highest and lowest observations. *N* = 3. **3C:** Plots showing division time of *NEUROG3*+ cells. The upper panel shows the division times with respect to the start of the movie (real time), and the lower panel shows division times normalized to the onset of GFP in each cell (relative time). Colors indicate independent experiments. *N* = 3, *n* =27. **3D:** Representative fluorescence tracing plots for three selected dividing *NEUROG3*+cells, showing GFP and RFP fluorescence intensities over time for different division modes. Dark and light lines indicate daughter cells after the division event. Fluorescence intensities were scaled between 0 and 1. The right panel shows the quantification of different division modes. *N* = 3, *n* = 29. **3E:** Plots showing fluorescent traces of all traced cells together for dividing (until division point) and non-dividing *NEUROG3*+ cells. On the right panel, fluorescence intensities were scaled between 0 and 1. Dark green, light green, dark red, and light red lines show smoothened mean intensities. The distribution represents individual values for each track per time point and the gray area shows the confidence interval. *N* = 3, *n* = 52 for nondividing cells and *N* = 3, *n* = 27 for dividing cells. **3F:** Flow cytometry-based live cell cycle analysis of GFP-low and GFP+ cells from different stages of *in vitro* differentiation. Bars indicate mean percentage values +-SEM. *N* = 2 for S3D2 and *N* = 4 for all the other groups (extended plot provided in S3D).

In 80% of the division events that we could record and categorize, both daughters of a GFP+ cell had similar trajectories of fluorescence accumulation, which we qualify as a symmetric division. For the remaining 20%, we observed that one of the daughter cells continued to upregulate GFP and RFP, whereas the other stayed at a low fluorescence intensity in both channels, which we termed as an asymmetric division (Figure 3D). This observation suggests that either *NEUROG3*+ cells act as an endocrine progenitor pool in which asymmetric cell division ensures the maintenance of a proliferative population with low *NEUROG3* levels (Bechard et al., 2016), or that at this early point of *NEUROG3* transcription, the cells can still revert, gradually losing their fluorescent signal. However, no cells were seen to divide more than once over 5 days of imaging. To determine the fate of daughter cells generated through asymmetric division and to assess the long-term proliferative capacity of the dividing cells, a longer tracing of the daughter cells would be required. Due to the crowding and mounding of the GFP+ cells at subsequent days of the differentiation protocol, our 2D imaging system was insufficient in answering these questions quantitatively. Therefore, we approached this question by utilizing a 3D culture system that allows long-term tracking and avoids confluency, as detailed in a subsequent section.

Comparing the dividing and non-dividing *NEUROG3*+ cells revealed that the dividing *NEUROG3*+ cells accumulated GFP and RFP intensities with similar slope and profile on average to their non-dividing counterparts (Figures 3E and S3B). As *Neurog3* is known to initiate an epithelial-to-mesenchymal-like program (Gouzi et al., 2011; Rukstalis and Habener, 2007), we also compared the movement speed between these two groups but did not find any significant differences in the average speed for the dividing and nondividing *NEUROG3*+ cells during the span of tracing (Figure S3C). Proliferating GFP+ cells were found throughout the differentiation protocol, analyzed through Stage 3 Day 2 to Stage 6 Day 1 using flow cytometry-based live cell cycle analysis as well as immunostaining at selected timepoints (Figures 3F and S3D-E). Even though the percentage of GFP+ cells in the G2M phase decreased over time due to the accumulation of GFP+ cells further along the differentiation path in culture, the percentage of cells in the G2M phase was relatively stable over time within the GFP low compartment, further demonstrating that the newly-induced *NEUROG3*+ cells with low transcription levels are more likely to divide (Figures 3F and S3D). Their percentage of cells in the cell cycle was similar to those found in GFP-pancreas progenitors. Taken together, we show that human endocrine progenitors can divide after *NEUROG3* onset once, mainly in the early phase of *NEUROG3* expression.

### Newly-emerging human endocrine progenitors differentiate in a gradual and uniform trajectory

The increased propensity to proliferate at low *NEUROG3* levels suggested that there may be molecularly and functionally different populations among *NEUROG3*+ cells. To compare their molecular identity globally, we next performed deep single-cell RNA sequencing (scRNA-seq) using the plate-based Smart-seq2 platform (Picelli et al., 2013), prioritizing sequencing depth over the number of cells analyzed. During our 2D *in vitro* human pancreatic differentiation, we index-sorted pancreatic cells by fluorescence-activated cell sorting (FACS) with no GFP, low GFP, and high GFP levels in predetermined ratios at Stage 4 Day 1, corresponding to the end of the live imaging tracks (Figures 1C and 4A). Reporter efficiency based on the FACS-based GFP categories of *NEUROG3* mRNA-containing cells from the scRNA-seq was calculated to be 97% (Figure S4A). Unsupervised cell embedding using uniform manifold approximation and projection (UMAP) revealed that cells align with their FACS-based GFP categories, reflecting their differentiation trajectory (Figures 4B, 4C, and S4B). Most cells in the G2M phase of the cell cycle formed a proliferative cluster, which included pancreatic progenitors as well as some *NEUROG3* mRNA-containing cells, further supporting that some of the *NEUROG3*+ cells do indeed express the transcriptional signature of dividing cells (Figures 4C and 4D). However, a few proliferative cells were detected all along the differentiation trajectory, confirming the observations from the imaging.

**Figure 4:**
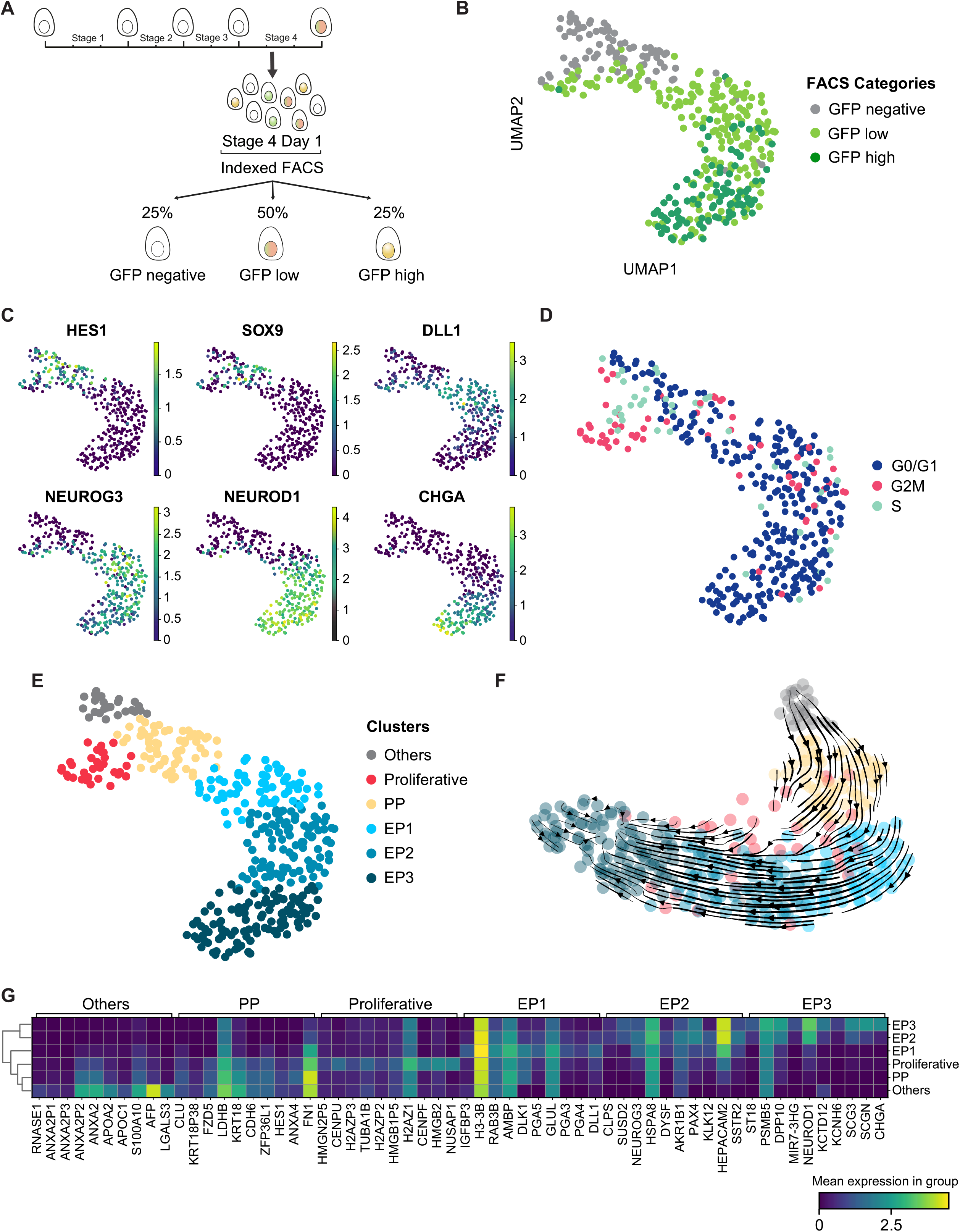
Deep single-cell sequencing of *NEUROG3*+ cells during human pancreatic endocrine differentiation indicates a gradual differentiation trajectory prior to hormonal transcription. **4A:** Schematic illustration depicting the indexed FACS strategy for the single-cell RNA-seq experiment, indicating the percentage of cells in different GFP categories sequenced. **4B:** UMAP colored by FACS-based GFP categories. **4C:** Selected normalized marker gene expression levels projected on the UMAP, highlighting pancreatic differentiation trajectory. **4D:** UMAP colored by sequencing-based cell cycle categories. **4E:** UMAP colored by detected clusters. **4F:** RNA velocity projected onto the UMAP produced after cell cycle regression, colored by detected clusters. Arrows indicate the extrapolated state. **4G:** Heatmap plot demonstrating the expression level of top 10 marker genes per cluster. PP: Pancreatic Progenitors, EP: Endocrine Progenitors.

The results indicated an overall differentiation continuum towards a pancreatic endocrine fate. When we performed clustering analysis on our dataset, we categorized the cells as pancreatic progenitors (PP), proliferative cells, and cells in the early steps of the endocrine differentiation path, named EP1, EP2, and EP3 (Figures 4E-G and S4C). We note that cells with low GFP expression are detected among cells with higher expression in EP2 and EP3, suggesting that cells with lower GFP expression activate differentiation similarly to cells with high levels (Figure 4A). Finally, we identified a cluster of cells expressing early liver marker genes such as alpha fetoprotein (*AFP*), which were also observed in other pancreatic differentiation datasets (Gonçalves et al., 2021; Veres et al., 2019) and assumed to be an earlier progenitor population (Figures 4E-G, and S4C). Cell cycle categories for each identified cluster are provided in Figure S4D, showing the decrease in the percentage of cycling cells from PP to EP3. We also noted the expression of known cell cycle exit-related genes in our data, such as *CDKN1C*, *BTG2*, and *GADD45A* (Figure S4E).

**Figure 5:**
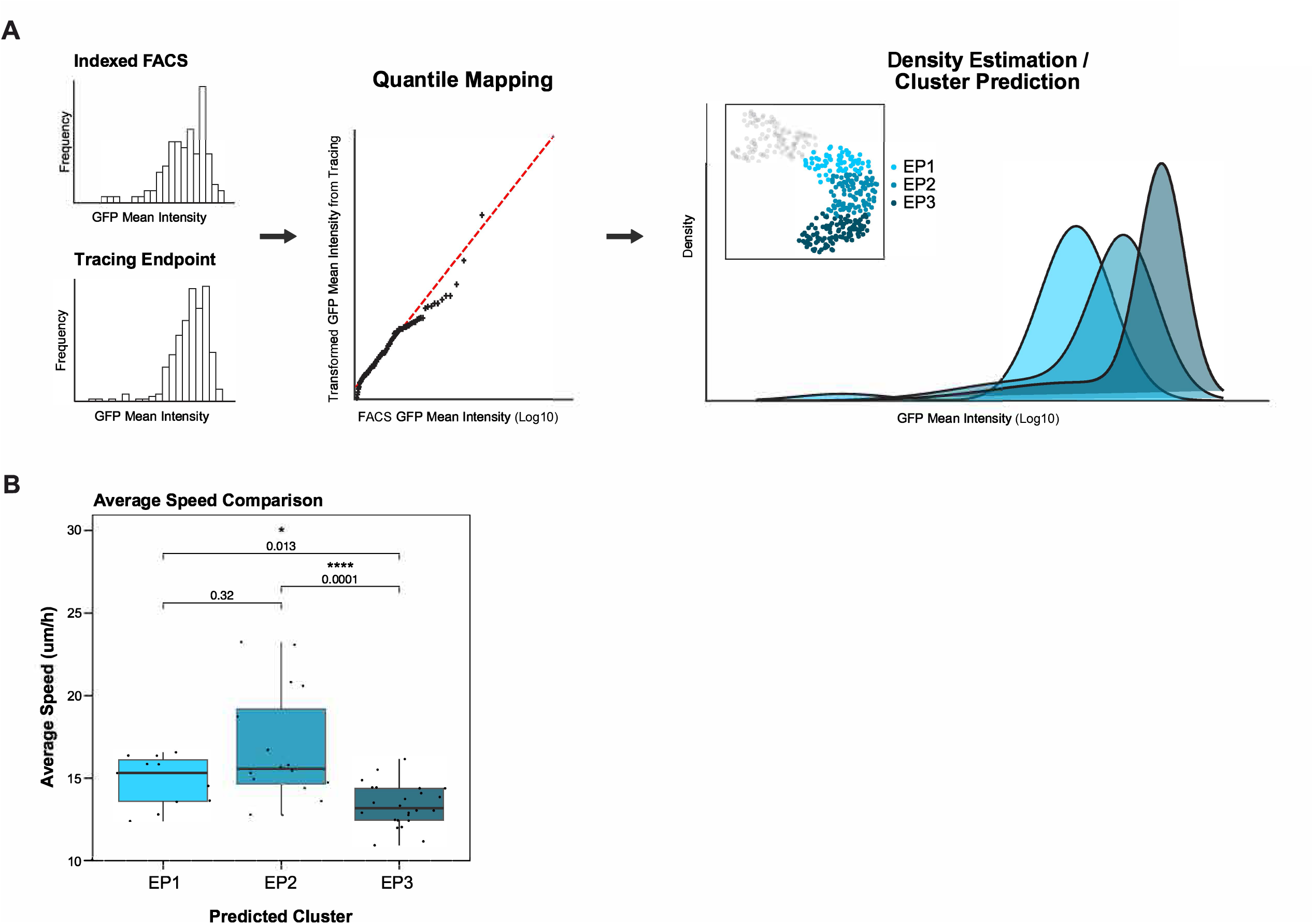
Correlating live imaging and single-cell sequencing of early hESC-derived endocrine progenitors uncovers differences in physical properties in cells with different *NEUROG3* levels. **5A:** Schematic outline of the analysis pipeline. **5B:** Comparison of average movement speeds of tracked cells according to their predicted EP clusters. P-value was determined by the Kruskal-Wallis test. *N* = 3, *n* = 52.

RNA velocity analysis performed on the data after cell cycle regression further supported our earlier conclusion, indicating a high degree of transcriptional changes, with a faster change in transcription at the onset of *NEUROG3* expression (Figure 4F). In addition, proliferative *NEUROG3*+ cells without cell cycle signature were mostly scattered to EP1 and EP2 clusters, indicating their transitioning status with fast progression towards the endocrine lineage (Figure 4F). At this point in endocrine differentiation at Stage 4 Day 1, we did not yet find many hormonal transcript-expressing cells or cells expressing marker genes for specific endocrine subtypes, such as aristaless related homeobox (*ARX*),indicating their further endocrine fate (Figure S4B). *CHGA*+ cells were the most mature population in our dataset, suggesting the transition from *NEUROG3* onset to hormone transcript expression takes more than 26 hours (Figure 4C). An exception is Ghrelin (*GHRL*) (Figure S4B), which has previously been shown to be activated early in endocrine progenitors *in vitro* and *in vivo* (Petersen et al., 2017; Ramond et al., 2018).

### Mapping live imaging to single-cell RNA sequencing enables correlating cellular dynamics to transcriptomics

Our scRNA-seq timepoint matches with the endpoint of tracing from the live imaging experiments. Using quantile transformation, we aligned relative GFP intensity values from the indexed FACS prior to scRNA-seq with those at the last time point of tracing from the live imaging movies (Figure 5A). Using Gaussian mixture modeling, we statistically predicted which sequencing cluster each traced non-dividing *NEUROG3*+ (GFP+) cell from live imaging would fall into (Figure S5A). With this mapping strategy, it is possible to correlate the physical properties of cells as observed by live image analysis to their transcriptomic profiles (Figure S5B). Building upon prior literature on the motility of *NEUROG3*+ cells (Gouzi et al., 2011), we compared the average moving speed of the cells in these three predicted clusters with different levels of *NEUROG3* expression, and we observed that EP3 cells were significantly slower on average than EP1 and EP2 cells (Figure 5B and S5C). We did not find a bias in the location of the cells predicted to belong to different transcriptomic clusters (Figure S5C). Taken together, our method of statistical mapping transcriptomic clusters to dynamic behaviors of cells in live imaging uncovered decreased motility in human pancreatic endocrine progenitors as the cells decrease NEUROG3.

### *NEUROG3*+ human pancreatic cells behave similarly in 2D and 3D culturing conditions

To validate our findings on human *NEUROG3*+ cells from 2D adherent cultures, next, we utilized the 3D human pancreatic progenitor spheroid culture system we previously developed (Gonçalves et al., 2021). In this system, hESCs or hiPSCs are differentiated until the pancreatic progenitor stage, and then, the cells are dissociated and seeded in a Matrigel dome with a minimal component culture medium to maintain and expand the pancreatic progenitors. By default, about 1-2% of cells spontaneously differentiated into endocrine cells in this medium. We thus induced endocrine differentiation in pancreatic progenitors 48 hours after seeding by adding a gamma-secretase inhibitor XX to the culture medium for downregulation of hes family bHLH transcription factor 1 (*HES1*), an important Notch signaling effector in the pancreas that directly binds to *NEUROG3* promoter and suppresses its transcription (Jensen et al., 2000; de Lichtenberg et al., 2018). We started the live imaging experiment using light-sheet microscopy 48 hours after treatment (Figure 6A). We detected both GFP and RFP signals within the differentiating spheroids (Figure 6B), and a number of GFP+ cells were also observed to divide in this system (Figure 6C). Similar to the 2D conditions, there was a high degree of heterogeneity in the *NEUROG3* expression levels and dynamics in the 3D human pancreatic cells (Figures 6D and S6A-B). Keeping this observation in mind, we quantified the expression period of the human NEUROG3 fusion protein using RFP as a proxy in single cells and found it to be around 40 hours (Figure 6D). The average GFP and RFP fluorescence accumulation rate and dynamics of the human NEUROG3 fusion protein within single cells were found to be very similar, with ramping up of *NEUROG3* transcription taking around 22 hours and the human NEUROG3 fusion protein taking around 11 hours even though the culturing system and the *NEUROG3* induction method were different (Figure 6E). This finding suggests that *NEUROG3* transcription kinetics may be invariable between these two different differentiation systems, and as long as there is a sufficient trigger for the initiation of promoter activation, *NEUROG3* transcription may occur with similar dynamics.

**Figure 6:**
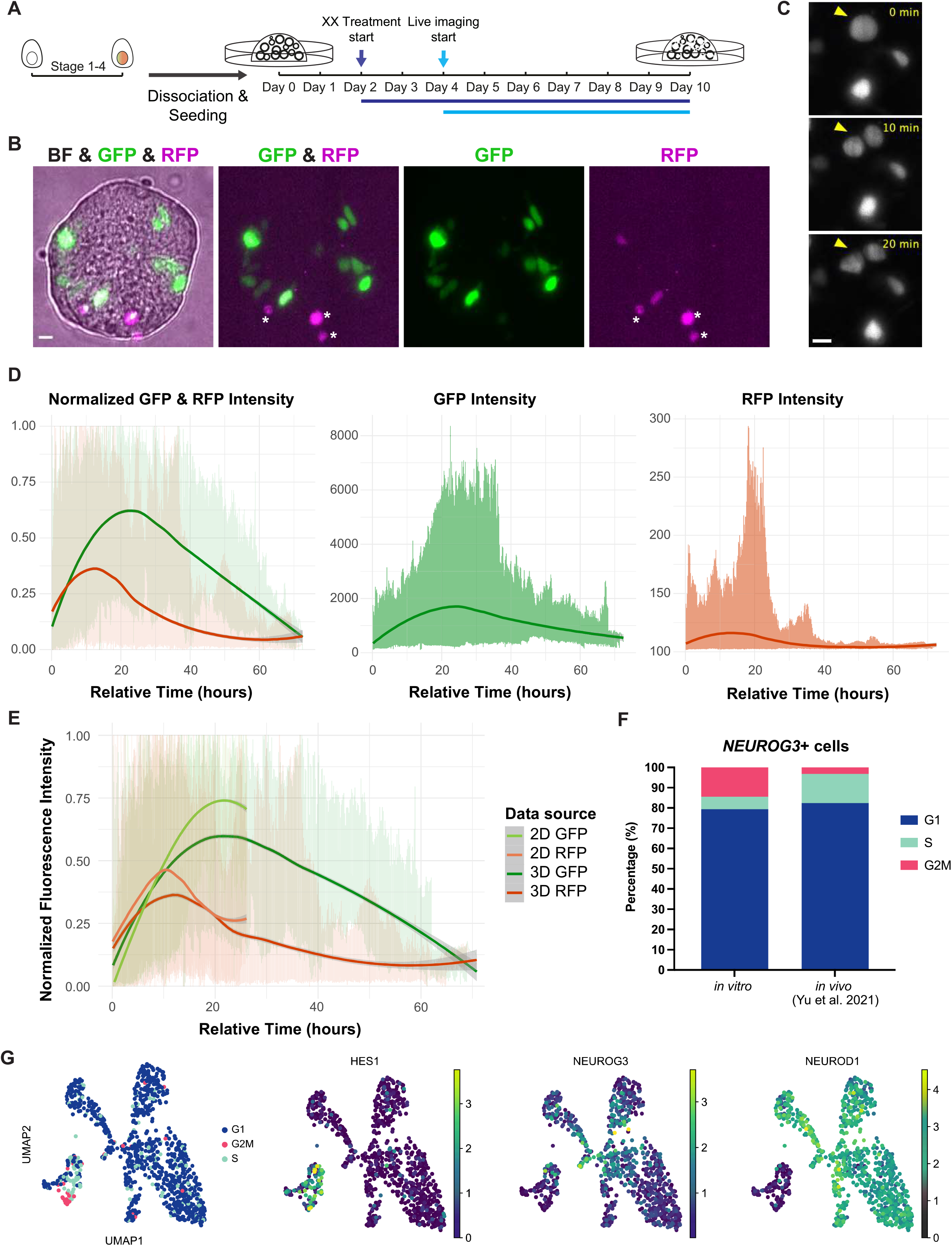
*NEUROG3* transcription and protein dynamics are very similar in 2D and 3D culture systems, and dividing *NEUROG3*+ cells exist in *in vitro* and *in vivo*. **6A:** Schematic representation of pancreatic progenitor spheroid culture and differentiation induction, as well as live imaging timeline. **6B:** Sample fluorescence microscopy image from differentiated spheroids at day 6 of culture, showing brightfield image (gray), GFP fluorescence (green), and RFP fluorescence (magenta). Asterisks indicate debris. Scale bars, 10 μm. **6C:** An example of a division event detected in a GFP+ cell during the live imaging of differentiating spheroids. Arrowheads indicate dividing and divided cells over time, with 3 panels showing 0, 10, and 20 minutes. **6D:** Plots showing fluorescent traces of all cells traced from the 3D culture. On the left panel, fluorescence intensities were scaled between 0 and 1. Dark green and dark red lines indicate smoothened mean fluorescence intensities. The distribution represents individual values for each track per time point and the gray area shows the confidence interval. *N* = 1, *n* = 30. **6E:** Comparison of the GFP and RFP intensities of traced cells from 2D and 3D human cell cultures. Light green and red lines and dark green and red lines indicate smoothened mean fluorescence intensities from 2D and 3D, respectively. The distribution represents individual values for each track per time point and the gray area shows the confidence interval. **6F:** Quantification of *NEUROG3*+ cells from scRNA-seq (*in vitro*) and Yu et al. 2021 (*in vivo*) according to their sequencing-based cell cycle categorization. 193 cells are represented for *in vitro*, and 969 cells are represented for *in vivo*. **6G:** UMAP plots generated from *NEUROG3*+ cells from Yu et al. 2021. The left panel indicates sequencing-based cell cycle categories, and the right panels show selected normalized gene expression levels projected onto the UMAP. Scale bars: 10 μm, BF: Brightfield.

Even though validating our results *in vivo* is not trivial, we compared some of our findings to the recent study by Yu et al., where different populations from human fetal pancreata spanning from 9 to 19 weeks post conception were sequenced using a deep single-cell RNA sequencing method (Yu et al., 2021). We found similar transcriptional signatures of EP1, EP2, and EP3 clusters to the reported human fetal pancreas-derived endocrine progenitor clusters (Table 1). Furthermore, scRNA-seq-based cell cycle categorization of *NEUROG3*+ human fetal pancreatic cells from Yu et al. revealed that some of these cells also express the transcriptional signature of cells that divide (Figure 6F and 6G), even though this dataset predominantly includes more mature endocrine populations than described here (Figure 6G). Therefore, this characteristic of early *NEUROG3*+ cells is reflected *in vivo* as well.

**Table 1 (Related to Figure 6):** Comparison of cell type-featured genes from the clusters of mSTRT-seq data of human fetal endocrine cells from Yu et al. 2021 to the top marker genes per detected cluster from this study.

|  | EP3 | EP2 | EP1 | PP | Cycling | Others | Color Code |
| --- | --- | --- | --- | --- | --- | --- | --- |
| 1 | CHGA | SSTR2 | DLL1 | FN1 | NUSAP1 | LGALS3 | Yu 2021 EP1 |
| 2 | SCGN | HEPACAM2 | PGA4 | ANXA4 | HMGB2 | AFP | Yu 2021 EP2 |
| 3 | SCG3 | KLK12 | PGA3 | HES1 | CENPF | S100A10 | Yu 2021 EP3 |
| 4 | KCNH6 | PAX4 | GLUL | ZFP36L1 | H2AZ1 | APOC1 | Yu 2021 EP4 |
| 5 | KCTD12 | AKR1B1 | PGA5 | CDH6 | HMGB1P5 | APOA2 | Yu 2021 ε |
| 6 | NEUROD1 | DYSF | DLK1 | KRT18 | H2AZP2 | ANXA2 | Yu 2021 δ |
| 7 | MIR7-3HG | HSPA8 | AMBP | LDHB | TUBA1B | ANXA2P2 | Yu 2021 β |
| 8 | DPP10 | NEUROG3 | RAB3B | FZD5 | H2AZP3 | ANXA2P3 | Yu 2021 α |
| 9 | PSMB5 | SUSD2 | H3-3B | KRT18P38 | CENPU | ANXA2P1 | Yu 2021 α/PP-Pro |
| 10 | ST18 | CLPS | IGFBP3 | CLU | HMGN2P5 | RNASE1 | Yu 2021 PP |
| 11 | CALB2 | GHRL | BTG2 | SERPINA1 | HMGN2P3 | VIM | not found |
| 12 | GCA | RASA4CP | SEMA6A | LDHBP2 | TPX2 | FTL |  |
| 13 | RGS4 | HSPA12A | H3-5 | SOX9 | TOP2A | LDHB |  |
| 14 | C22orf42 | PRICKLE2 | GADD45A | KRT18P15 | DLGAP5 | S100A16 |  |
| 15 | ESRRG | SCG2 | NREP | FTH1 | MAD2L1 | APOE |  |
| 16 | RASSF6 | ENC1 | SOX4 | FTH1P7 | PTTG1 | FTLP3 |  |
| 17 | MAP1B | AKR1B1P2 | FLRT2 | KRT8P3 | HMGB1P6 | FTH1P7 |  |
| 18 | OR51E1 | HSPA8P5 | FRZB | KRT18P17 | CKS1B | LDHBP2 |  |
| 19 | CDKAL1 | CADPS | TSC22D1 | KRT18P11 | HMGB1 | SAT1 |  |
| 20 | AMIGO2 | DEPP1 | ARF5 | ID3 | BIRC5 | ANXA3 |  |
| 21 | CPE | DCC | GLULP4 | ANGPT2 | TUBAP2 | GSTO1 |  |
| 22 | SCG2 | SEC61B | NPY | IFITM3 | HMGB1P1 | CA2 |  |
| 23 | PAM | HSPA8P1 | SPINT2 | KRT18P16 | KIF20B | SPART |  |
| 24 | CACNA2D1 | TAGLN3 | HNRNPA1 | KRT8 | AL132777.1 | OCLN |  |
| 25 | THSD4 | AKR1B1P3 | CTNNAL1 | B2M | HMGN2 | LY6E |  |
| 26 | KCNK16 | OLFML2B | SLC2A1 | KRT8P10 | RRM2 | TMSB10 |  |
| 27 | SYT4 | AL450311.1 | CELA3A | KRT18P8 | MKI67 | FTH1 |  |
| 28 | RAB26 | AP003108.1 | LINC02381 | KRT18P10 | CCNB1 | TTR |  |
| 29 | POLR2L | SEC11C | GSTA2 | STAT1 | UBE2C | FILIP1L |  |
| 30 | KCNK17 | MUC15 | HNRNPA1P48 | ANXA2 | KPNA2 | KRT18 |  |

### Dividing *NEUROG3*+ cells do not have a unique transcriptomic signature and are in transition to the endocrine progenitor lineage

Our analysis so far indicated that although dividing *NEUROG3*+ cells exist throughout differentiation and in different culture systems, they may neither have unique characteristics nor be a part of a stable population. To rule out the possibility that the limited sample size did not allow us to uncover transcriptional differences of the dividing *NEUROG3*+ cells, we also performed bulk RNA sequencing with four sorted sample populations: GFP+ G0/G1 cells, GFP+ G2M cells, GFP-G0/G1 cells, and GFP-G2M cells. Principal component analysis (PCA) revealed that GFP+ G2M cells are transcriptionally more similar to the GFP-cells, representing a transitory state along the path of endocrine differentiation (Figure 7A). Gene Set Enrichment Analysis (GSEA) showed, unsurprisingly, genes downregulated by KRAS activation and pancreas beta cell-related genes as being enriched in the GFP+ cells compared to GFP-cells (Figure 7B). Within the GFP+ G2M compartment, pancreas progenitor-related genes were downregulated, while endocrine progenitor-related genes were upregulated, with no unique genes or pathways specifically activated or inhibited that could be uncovered (Figure 7C). When cycling and non-cycling GFP+ cells were compared, the GSEA included genes related to G2M checkpoint and E2F targets, as expected from the cell cycle category, as well as genes upregulated by KRAS activation, Notch signaling-related, and epithelial-mesenchymal transition-related genes being enriched in the cycling GFP+ cells (Figures 7D and S7A). Supporting the idea that the cycling GFP+ cells have newly activated endocrine differentiation, the non-cycling GFP+ cells expressed more beta cell-related genes (Figures 7D and S7A). Taken together, dividing *NEUROG3*+ cells are in a transitional state towards endocrine differentiation, with differentiation initiation occurring before the cell cycle completion of pancreatic progenitors.

**Figure 7:**
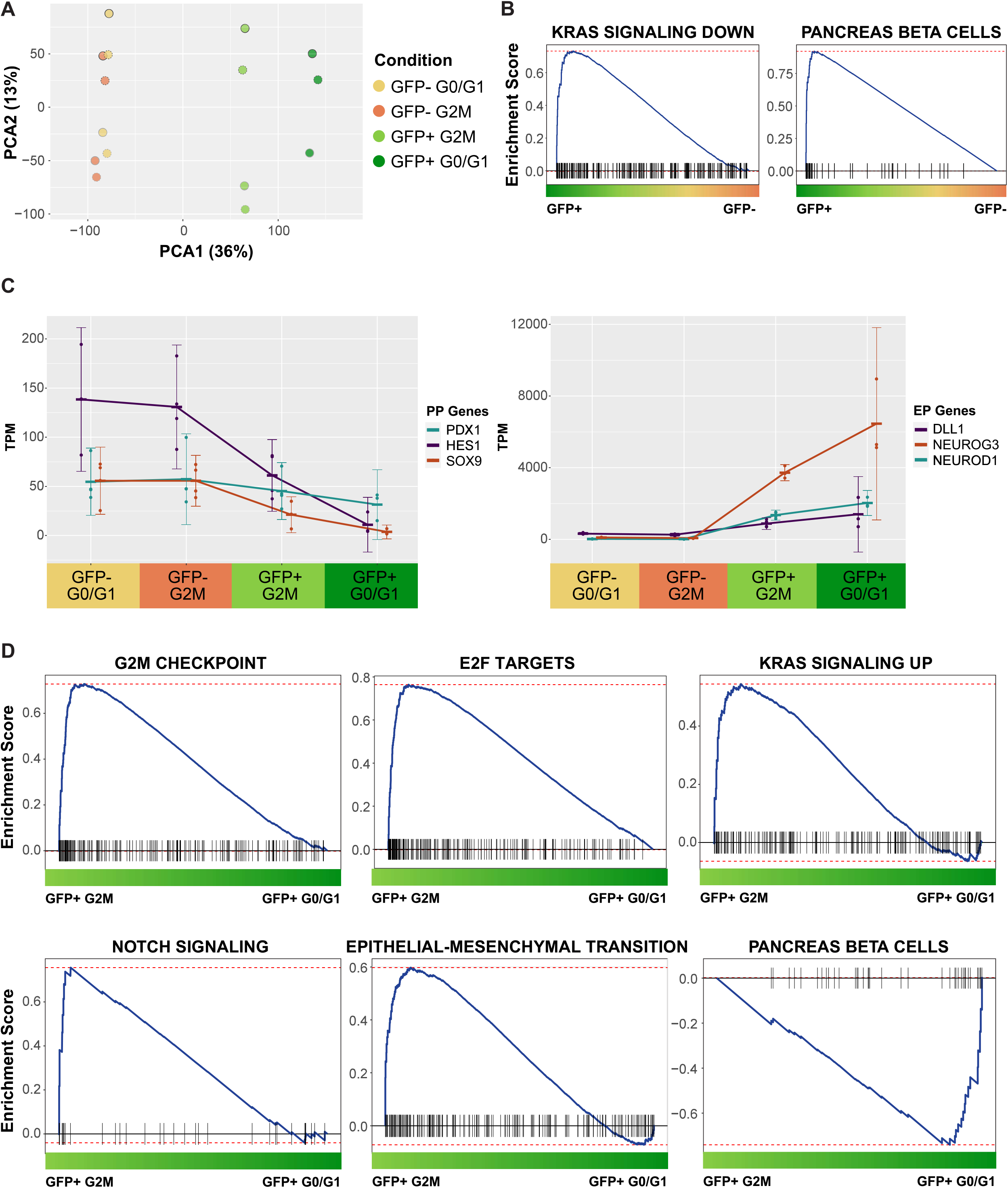
Dividing *NEUROG3*+ cells do not have a unique transcriptomic signature but are in transition to the endocrine progenitor lineage. **7A:** PCA plot showing the distance between the transcriptomes of GFP+ and GFP-G0/G1 and G2M cells. *N* = 3 for GFP+ G0/G1 and *N* = 4 for the other categories. **7B:** Gene Set Enrichment Analysis (GSEA) of GFP+ cells compared with GFP-cells. **7C:** Mean TPMs (Transcripts per Million) with standard deviation error bars for selected pancreatic progenitor genes (*PDX1*, *HES1*, and *SOX9*) and endocrine progenitor genes (*DLL1*, *NEUROG3*, and *NEUROD1*) per sequencing category. **7D**: Selected GSEA panels of GFP+ G2M cells compared with GFP+ G0/G1 cells (full list provided in S7A).

## Discussion

### *NEUROG3* dynamics are slower in humans compared to mice

The human pancreas develops over a longer period but is also larger than in mice, with its ratio of endocrine cells being similar. Using the GFP signal from our reporter as a proxy for transcription, we show that in the human pancreatic cells, *NEUROG3* transcription reached its peak in around 22 hours, whereas in a mouse line where *Neurog3* promoter was used to drive RFP expression (Kim et al., 2015), it peaked at around 12 hours. EGFP and TagRFP-T have similar folding rates (Shaner et al., 2008); therefore, the two-fold difference between human and mouse reporter expression is likely due to the transcription and reporter translation differences between species. Recent work in other systems such as the presomitic mesoderm and motor neurons showed that differences in global biochemical rates play a substantial role in creating the divergence in the timespan of development and that human cells display twice slower kinetics of protein expression and decay compared to the mouse cells (Matsuda et al., 2020; Rayon et al., 2020). Although we cannot directly compare the NEUROG3 protein expression length of individual human and mouse pancreatic cells, we observed that the accumulation of transcripts and protein were synchronous in the ascending phase, which infers that the protein is translated with no delay, suggesting that the protein accumulation is likely to be slower in human. Though a mouse Neurog3 fusion protein was recently generated, its dynamics have not yet been studied by live imaging (Bastidas-Ponce et al., 2019). In humans, NEUROG3 protein peaks 11 hours after its expression onset and remains for about 40 hours. The time it takes in our system to observe NEUROG3-RFP levels to reduce by half combines the protein decay and the progressively slower rate of new transcription after about 12 hours. The protein degradation measurements indicate that NEUROG3 half-life is shorter in mice, ranging from 12 to 15 minutes (Azzarelli et al., 2017; Roark et al., 2012) than in humans, 30-67 minutes (Zhang et al., 2019), though caution is needed as it was measured in less physiological systems, and the half-life was cell type-dependent (Zhang et al., 2019).

Though it is risky to claim the relevance of *in vitro* observations to the development in the womb, we show robustness in the differentiation timing across systems. Induction of *NEUROG3* expression in a 3D pancreatic progenitor culture system (Gonçalves et al., 2021) showed that human *NEUROG3* transcription and protein dynamics are very similar between different culture systems (2D vs. 3D), media with different growth factors (high bFGF in 3D) and even different induction methods (Notch inhibition in 3D, FGF7, retinoic acid, SANT1, LDN and TPB in 2D); suggesting that these measurements may be relevant to an *in vivo* setting.

An important new observation we made was the high degree of variability in the *NEUROG3* peak expression among cells, both in the 2D and the 3D conditions. It is thus likely that endocrine differentiation occurs above a low threshold of *NEUROG3* expression, and some cells vastly exceed this threshold. A pending question is whether all these cells give rise to endocrine cells. In mice, previous experiments using a BAC reporter for *Neurog3* traced some progeny to the exocrine lineage (Cortijo et al., 2012; Schonhoff et al., 2004). Although we could show that our cultures lead to the production of numerous hormone-expressing cells as they mature during *in vitro* pancreatic differentiation, tracing single cells until this maturity level was not possible due to the technical limitations of image analysis in 2D. Further work on the 3D pancreatic culture and differentiation platform can enable us to overcome these limitations. However, our single-cell RNA sequencing experiments do not show multiple trajectories downstream of *NEUROG3* expression, as one would assume to observe if some populations of *NEUROG3*+ progeny had taken an exocrine fate or reverted to pancreas progenitors. We observed that after human pancreatic cells initiate *NEUROG3* transcription, they undergo a uniform transcriptional change until they initiate *CHGA* transcription and that many GFP low cells mingle with the GFP high cells in the progression to EP2 and EP3. It remains possible that endocrine progenitors can revert after this step.

### *NEUROG3*+ human pancreatic cells can divide but do not act as a specialized population

In this work, we showed that *NEUROG3*+ human pancreatic cells could divide in different culture systems based on live imaging, flow cytometry-based cell cycle analysis, as well as RNA-seq-based cell cycle categorization. Dividing *NEUROG3*+ cells exist throughout the *in vitro* human pancreatic differentiation, and the cells are more likely to divide when they have low *NEUROG3* levels. At these early steps, they do so almost as effectively as pancreas progenitors. Following the daughters after division through live imaging, we saw that different division modes exist, with some daughter pairs both upregulating *NEUROG3* (80% of the cases), and others where only one daughter upregulating *NEUROG3* while the other staying low (20% of the cases). It is currently unclear whether the asymmetric division results in one daughter turning *NEUROG3* expression off and reverting to a pancreatic progenitor fate or sustaining a low level of *NEUROG3* transcription.

Even though human pancreatic cells can divide after *NEUROG3* initiation, we could neither detect any cell dividing more than once, even in 5-day movies, nor find any dynamic or transcriptomic signature associated with these cells to indicate they are a specialized population. Prior to division, these cells exhibited the same *NEUROG3* transcription and protein dynamics as their non-dividing counterparts, and no behavioral differences were recorded between them by live imaging. Bulk RNA sequencing of endocrine progenitors in G2/M only uncovered cell division signatures and a signature of a transitionary state between pancreas progenitors and endocrine progenitors. Together with the observation that cell cycle exit-related genes such as *CDKN1C* (de Lichtenberg et al., 2018), *GADD45A*, and *BTG2* (Krentz et al., 2018; Schreiber et al., 2021) turn on in the EP1 population with *NEUROG3*, we conclude that *NEUROG3*+ human pancreatic cells do not act as a stable endocrine progenitor pool as proposed in mice (Bechard et al., 2016; Larsen et al., 2017). Rather, these cells most likely finish their last cell cycle round before further *NEUROG3* upregulation and cell cycle exit. In mice, we had observed a slightly lower occurrence of *NEUROG3*+ cell division, at around 3.2% (Kim et al., 2015). The higher percentage of dividing *NEUROG3*+ cells in humans may be a specific human property or could mean that the sensitivity of our dual human *NEUROG3* reporter line allows us to discern the low *NEUROG3*-expressing cells better. Recent mouse studies explored the induction of NEUROG3 and found that G1 lengthening in trunk progenitors drives endocrine differentiation by slowing NEUROG3 phosphorylation, thereby increasing the stability of mouse NEUROG3 protein (Azzarelli et al., 2017; Krentz et al., 2017; Roark et al., 2012). Even though phosphorylation of human NEUROG3 was also shown (Krentz et al., 2017), the phosphorylation sites and mechanisms are most likely not the same in humans (Zhang et al., 2019). If phosphorylation in humans acts mechanistically similarly to what was proposed in mice, our observations suggest that it is unlikely to be in cells that would remain in a cycling *NEUROG3-low* state with recurrent degradation over multiple cell cycles, but this may participate in a feed-forward loop enabling the ramping up of NEUROG3. The role of phosphorylation could be further investigated with our reporter by perturbing the phosphorylation of NEUROG3 and investigating its effect on *NEUROG3* dynamics and proliferation.

### Initial steps after *NEUROG3* induction are similar *in vitro* and *in vivo*,and *NEUROG3*+ cells from human fetal pancreata can also divide

A limitation of work on *in vitro* systems addressing human development is the difficulty of assessing their relevance to the *in vivo* context (Gonçalves et al., 2021). Our single-cell RNA sequencing dataset indicated that human pancreatic cells undergo a gradual differentiation process after *NEUROG3* initiation. The similarity between *in vitro* and *in vivo* gene expression profiles indicates that the downstream transcriptional program after *NEUROG3* expression is robust and can be captured faithfully in human *in vitro* pancreatic differentiation systems (Yu et al., 2021). This finding extends previous observations as the convergence of the transcriptional program during endocrine induction was also reported in less extensive qPCR-based single-cell profiling of *in vivo* and *in vitro*-derived pancreatic cells (Ramond et al., 2018).

Even though the dataset from Yu et al. predominantly included more mature endocrine populations than described here, finding *NEUROG3*+ human fetal pancreatic cells with a proliferation signature *in vivo* further validates the relevance of this study to the human fetal endocrine pancreas development. The similar transitions in gene expression profiles found in their pseudotime analyses *in vivo* and our analyses *in vitro* also suggest a similar differentiation progression.

### Mapping live imaging to single-cell RNA sequencing connects dynamic behaviors to the transcriptomic signature

Linking single-cell sequencing profiles to functional readouts is a new emerging challenge in cell omics. In this study, by using a plate-based RNA-sequencing platform in combination with indexed FACS, we were able to recall fluorescence intensities of sequenced cells in different transcriptomic clusters to statistically correlate them with the intensities of the live-imaged *NEUROG3*+ cells. With this mapping strategy, it is possible to link the physical properties of any chosen cell in the movies, such as the movement speed exemplified here, to their transcriptomic profiles in a statistical way. Using this approach, we found that the most mature endocrine progenitor cells with the highest *NEUROG3* levels in our analysis, EP3 cells, start to slow down in migration compared to EP1 and EP2 cells. We had previously shown that NEUROG3 triggers motility and partial epithelial delamination (Gouzi et al., 2011), which we confirmed in our 3D system here in humans at endogenous expression levels, though under endogenous conditions, the *Neurog3*+ cells and their endocrine progeny remain connected to the epithelium (Sharon et al., 2019). Our observation extends these findings by suggesting that the motility of cells decreases after NEUROG3 downregulation starts, which occurs in the EP3 cells. We also identify specific candidates from the EP2 signature associating with the motility peak such as the serine protease KLK12 (Figure S4C). Such a methodology can be applied to other dynamic properties that can be extracted from live imaging in diverse biological systems to obtain a more comprehensive overview of a system in dynamically changing landscapes.

Taken together, the mapping method proposed could be transposed to other developmental processes to link changes in gene expression genome-wide to observable events in live imaging. Conceptually, our study reveals that differentiation dynamics during the initiation of endocrine commitment in human pancreas development is slower than in mice and is a noisy process with regards to expression levels but very reproducible in its timing between cells and between conditions. Of importance for therapeutic avenues, we show that while an endocrine-committed proliferative population would have been convenient to produce pure populations of human pancreatic endocrine cells in large numbers for cell therapy of diabetes mellitus, there is no evidence for sustained proliferation potential nor subpopulations of proliferative cells.

## Acknowledgments

The authors would like to thank the DanStem Genomics Platform, Stem Cell Culture Platform, Flow Cytometry Platform (especially Paul van Dieken), and Imaging Platform for training, technical expertise, support, and the use of instruments. Additionally, we thank the Scientific Computing Facility (especially Noreen Walker, Gayathri Nadar, and Andre Gohr), Organoid and Stem Cell Facility, FACS Facility, and Light Microscopy Facility at MPI-CBG for their support and technical help. The Deep Sequencing Facility of the CMCB Technology Platform (Center for Regenerative Therapy in Dresden) provided invaluable service with library preparation and sequencing for the RNA sequencing experiments. The Novo Nordisk Foundation Center for Stem Cell Biology is supported by a Novo Nordisk Foundation grant number NNF17CC0027852. B.S.B.-T. is supported by the Copenhagen Bioscience PhD program financed by the Novo Nordisk Fonden (NNF16CC0020994).

## Author Contributions

Conceptualization, B.S.B.-T., Y.H.K., and A.G.-B.; Methodology, B.S.B.-T., Y.H.K., L.H., F.L., C.Z., and A.G.-B.; Investigation, B.S.B.-T., and J.V.D.; Resources, A.G.-B.; Writing – Original Draft, B.S.B.-T., Y.H.K., and A.G.-B.; Writing – Review & Editing, B.S.B.-T., L.H., F.L., C.Z., Y.H.K., and A.G.-B.; Visualization, B.S.B.-T.; Supervision, Y.H.K. and A.G.-B.

## Methods

### Culturing of human embryonic stem cells

The H1 hESC line was purchased from WiCell. Approval to work on hESC was obtained from De Videnskabsetiske Komiteer, Region Hovedstadenunder number H-4-2013-057 and Central Ethics Commission for Stem Cell Research (ZES) and the Robert Koch Institute (RKI) (AZ: 3.04.02/0148). The hESCs were cultured on hESC-Qualified Matrigel (Corning) and maintained using mTESR1 medium (Stem Cell Technologies). The media was changed daily, and the cells were kept at 37°C and 5% CO2. Approximately every 3-4 days, after the cell confluence had passed 70-80%, the cells were passaged by dissociating with TrypLE (Thermo Fisher) and replating at a density of 40,000 cells/cm2. During the first 24 hours after passaging, hESCs were supplemented with 10 μM ROCK inhibitor Y-27632 (Sigma). The cells were periodically tested and cleared for mycoplasma and karyotypic abnormalities. The hESC work was conducted according to the guidelines.

### Construction of the dual *NEUROG3* reporter hESC line

For the construction of the dual *NEUROG3* reporter hESC line, insertion of GSx3 linker-TagRFP-T-P2A-EGFP-NLS-STOP coding sequence immediately before the stop codon of the *NEUROG3* coding sequence in H1 hESCs was achieved by homology-directed repair of a CRISPR/Cas9-induced DNA break.

A guide RNA targeting the *NEUROG3* locus was designed with CRISPR design (http://crispr.mit.edu), using the pX330 vector that encodes for human codon-optimized SpCas9 (Cong et al., 2013). The DNA plasmid included around 1000 base pairs-long homology arms from *NEUROG3* genomic DNA with a point mutation of PAM sequence in a vector backbone (pBluescriptSK), with the desired construct and a floxed selection cassette. The fragments were assembled using the Seamless cloning kit (Thermo Fisher), according to the manufacturer’s instructions. The two plasmids were co-electroporated into H1 hESCs, and antibiotic selection was performed. After the initial screening, surviving colonies were analyzed for correct integration. Following validation, the selection cassette was excised. From the two heterozygous reporter colonies that were validated, the choice was made on *in vitro* pancreatic differentiation efficiency.

The karyogram of the dual NEUROG3 reporter was analyzed on metaphase spreads at P14 (Cell Guidance Systems, Cambridge, UK) and P17 (Organisation Genetische Diagnostik Institut für Klinische Genetik Medizinische Fakultät Carl Gustav Carus TU Dresden).

### *In vitro* differentiation into the pancreatic lineage

*In vitro* differentiation of hESCs into pancreatic endocrine cells was performed as previously described (Petersen et al., 2017; Ramond et al., 2018), based on the protocol published by Rezania et al. (Rezania et al., 2014), with minor adjustments. In summary, hESCs were dissociated into single cells using TrypLE and resuspended in mTeSR1 with 10 μM ROCK inhibitor Y-27632. The cells were seeded at a density of 350,000 cells/cm2 onto 1:30 diluted growth-factor reduced Matrigel (Corning). The cells were incubated for 24 hours in mTeSR1 medium before transitioning to the differentiation media. Modifications from the original protocol (Rezania et al., 2014) include the usage of MCDB131 medium (Life Technologies) as the basal medium throughout differentiation instead of BLAR medium, activin A (PeproTech) at 100 ng/ml during stage 1 instead of GDF8, CHIR99201 (Axon Medchem) at 3 μM and 0.3 μM for the first two days of Stage 1 instead of MCX-928, and keeping the cells in 2D culture throughout the protocol instead of transitioning to air-liquid interface at Stage 5. The modified protocol’s differentiation efficiency was reported in Petersen et al. (Petersen et al., 2017) and is further documented for the reporter line in this paper (Figures 1 and S1). For the live imaging experiments, reporter hESCs and H1 hESCs were mixed in a 1:1 ratio and then differentiated for the ease of tracking fluorescent cells.

The generation of human pancreatic progenitor spheroids was performed as previously described in Gonçalves et al. (Gonçalves et al., 2021). Pancreatic progenitors differentiated from hESCs at the end of Stage 4 using the described pancreatic differentiation protocol were dissociated into single cells with TrypLE and resuspended in a 3:1 mixture of growth-factor reduced Matrigel and expansion medium (DMEM/F12 Glutamax (Thermo Fisher), 10 μM ROCK inhibitor Y-27632, 64 ng/ml FGF2 (Peprotech), and B-27 supplement (Thermo Fisher)). The cells were seeded at a density of 1000 cells/μl of Matrigel mixture onto pre-warmed Nunc cell culture treated 4-well dishes (Thermo Fisher) to form a 3D dome. The expansion medium was added after Matrigel polymerization and replenished every 3 days. The spheroids were passaged every 10 days. For the induction of endocrine differentiation in the pancreatic progenitor spheroids, 48 hours after seeding, 100 nM gamma-secretase inhibitor XX (EMD Millipore) was added to the expansion medium and was replenished every other day.

### Live imaging and cell tracking

Live imaging experiments with the 2D differentiation system were carried out using a Deltavision microscope (PerkinElmer) with the 20X objective in a humidified, heated, and CO2-controlled chamber. Tiled positions (4×4 tiles, 16 tiles per acquisition) were scanned every 6 minutes. At the end of the imaging, the cultures were immediately fixed with 4% PFA (Thermo Fisher) for immunocytochemistry. Imaris (Bitplane, Switzerland) software was used to perform manual tracking of NEUROG3+ cells over time, using spots of 5 μM diameter.

For the 3D cultures, a Viventis light-sheet microscope was used with a Nikon 25x 1.1 NA water immersion objective in a humidified, heated, and CO2-controlled chamber. Set positions were images every 10 minutes. Manual tracing of NEUROG3+ cells was performed with Imaris (Bitplane, Switzerland) software, using spots of 5 μM diameter.

### Flow Cytometric Analyses

For the flow cytometry-based live cell cycle analysis, at different time points during the human *in vitro* pancreatic differentiation protocol, the cells were dissociated and collected as described above and reconstituted in FACS buffer (1% BSA (Sigma) in 1x PBS (Thermo Fisher)). Ghost dye red 780 (Cell Signaling Technology) for live/dead stain and Hoechst 33342 Ready Flow Reagent (Thermo Fisher) were added with the concentration of 1 μl/1,000,000 cells and 2 drops/1,000,000 cells in 1 mL, respectively. The cells were incubated at 37°C for 1 hour, followed by filtering with a 35 μm cell strainer and flow analysis on a BD FACSAria (BD Biosciences).

For the fixed flow cytometric analyses, after dissociating cells to a single cell suspension, the cells were washed with 1x PBS and incubated on ice in the dark with 1 μl of live/dead stain (Ghost dye red 780) in 1x PBS for 10 minutes. Afterward, the cells were washed with 1x PBS and fixed with 4% PFA on ice for 20 minutes. After spinning down, the cells were resuspended in a permeabilization buffer (0.2% Triton X-100 (Sigma) and 5% Donkey serum (Sigma) in 1X PBS) and incubated on ice for 30 minutes. After pelleting, primary antibodies in a blocking buffer (0.1% Triton X-100 and 5% Donkey serum in 1X PBS) were added, and the samples were incubated at 4°C overnight. The next day, the cells were washed with an IC buffer (1% BSA in 1x PBS) and incubated with the secondary antibodies in a blocking buffer at room temperature for 45 minutes. Following 3 washes with the IC buffer, the cells were passed through a 35 μm cell strainer, and flow analysis was performed on a FACSAria or a Sony MA900 cell sorter (Sony). The primary antibodies for flow cytometry were: rat anti-C-peptide-Alexa Fluor 647 (1:200, 565831, BD Pharmigen) and mouse anti-Glucagon-BV421 (1:100, 565891, BD Pharmigen).

### Immunostaining

For the staining of hESC-derived cultures, the wells were washed with 1X PBS and fixed with 4% PFA at room temperature for 30 minutes. Afterward, the cells were washed with 1X PBS and permeabilized with 0.5% Triton X-100 in 1X PBS for 10 minutes. The cells were incubated in blocking buffer (0.1 M Tris-HCl (pH 7.5), 0.15 M NaCl, 0.5% TSA Blocking Reagent (PerkinElmer)) for 30 minutes, followed by overnight incubation at 4°C with primary antibodies diluted in 0.1% Triton X-100 in 1X PBS. The following day, cells were washed 3 times with 1X PBS for 5 minutes each and incubated with secondary antibodies in 0.1% Triton X-100 in 1X PBS for an hour at room temperature and imaged with a Leica Thunder microscope. The primary antibodies were: sheep anti-NEUROG3 (1:1000, AF3444, R&D Systems), chicken anti-GFP (1:1000, ab13970, Abcam), RAT anti-RFP (1:1000, 5F8, Chromotek), rabbit anti-MKI67 (1:500, ab16667, Abcam), and rabbit anti-SOX9 (1:2000, AB5535, Millipore).

### Image analysis

Image processing prior to manual cell tracking in 2D and 3D and quantification of the overlap of NEUROG3 immunostaining and the fluorescence from the reporter line was performed using the ImageJ distribution Fiji (v1.53c) (https://fiji.sc/) (Schindelin et al., 2012).

For the image processing prior to manual cell tracing in 2D, an automated script was used to perform the following steps: Stitching tile positions, drift correction, background subtraction, and downscaling.

For the image processing prior to manual cell tracing in 3D, cropping and downscaling were applied.

For the quantification of the overlap of NEUROG3 immunostaining and the fluorescence from the reporter line, an automated script was used to perform the following steps: creating an additional image for merged channels segmentation by StarDist (Schmidt et al., 2018) (https://imagej.net/plugins/stardist); filtering by size (8 μm); background subtraction (rolling = 50); masking positive cells from the merged image (intensity > image mean); identifying overlapping cells between channels.

### FACS, Single-cell RNA sequencing, and downstream analysis

At Stage 4 Day 1 of the differentiation protocol, the cells were dissociated into single cells with TrypLE and resuspended into a FACS buffer containing 0.5% BSA and 2 mM EDTA (Thermo Fisher) in 1X PBS. The cells were index-sorted according to their GFP expression (categorized into low GFP and high GFP based on the distribution of global GFP values) into 384-well plates on a BD FACSAria and snap-frozen. 397 sorted cells were processed using a modified version of the Smart-seq2 protocol (Picelli et al., 2013).

FastQC (http://www.bioinformatics.babraham.ac.uk/) was used to perform basic quality control of the resulting sequencing data. Trimming of low quality bases, Smartseq2 adapter sequences and poly(A)/poly(T) stretches was done with cutadapt (v2.6) (Kechin et al., 2017) and the following parameters: “-q 5 -m 20 -a smartseq1=AAGCAGTGGTATCA - a smartseq2=AACGCAGAGTGCAGTGC -b smartseq3=AACGCAGAGTGCAGTGC -a smartseq4=CTGTCTCTTATA -a polyA=AAAAAAAA -a polyT=TTTTTTTTTT -b smartseq5=GGTATCAACGCAGA --times 5”. Trimmed fragments were aligned to the human reference genome hg38 with support of the Ensembl 92 splice sites using the aligner gsnap (v2018-07-04) (Wu and Nacu, 2010). The sequence of the reporter *NEUROG3* fusion construct was included in the reference without the *NEUROG3* sequence to allow for mapping to the endogenous *NEUROG3* gene. Counts per gene and sample were obtained based on the overlap of the uniquely mapped fragments with the same Ensembl annotation and the included fusion construct sequence using featureCounts (v1.6.3) (Liao et al., 2014). After removing the three sequencing controls (positive, negative, and bulk controls) from the count matrix, the remaining 397 cells were checked for overall quality using the scater package v1.16.2 (McCarthy et al., 2017). Detected outlier cells were removed from the data set, and the remaining 380 cells were further analyzed with Scanpy v1.8.1 (Wolf et al., 2018) to perform clustering with the Leiden algorithm (Traag et al., 2019) and marker gene detection using the Wilcoxon ranksum test. Gene set enrichment analysis was run against MSigDB hallmark genes sets with more than 15 and less than 500 genes (msigdb v7.2.1) (Liberzon et al., 2015; Subramanian et al., 2005) using fgsea v1.8.0 (Korotkevich et al., 2021) by applying an adjusted p-value cutoff of 0.01. For this analysis, the ranking of input genes was done using the negative log10 of obtained p-values which were then additionally multiplicated with −1 in case of a negative log2 fold-change.

For RNA velocity analysis, spliced and unspliced transcripts were quantified with the smartseq2 command of velocyto v0.17.17 (La Manno et al., 2018) and served as input for further analysis with scVelo v0.2.3 (Bergen et al., 2020) using dynamical modeling. RNA velocity was projected upon a UMAP obtained with Scanpy after additional regression of cell cycle scores assigned based on cell cycle genes defined in Tirosh et al. (Tirosh et al., 2016).

### RNA extraction and sequencing of human pancreatic cell populations

Bulk RNA sequencing was performed on cells from 4 independent *in vitro* pancreatic differentiation experiments. The cells were FAC-sorted into RLT buffer (QIAGEN) with 1%β-mercaptoethanol (Sigma Aldrich) according to their GFP fluorescence intensity and cell cycle category with live cell cycle staining described above (Sony MA900 sorter). RNA was extracted from each replicate and population using the RNeasy Micro Kit (QIAGEN), according to the manufacturer’s guidelines. Subsequent RNA quality assessment was done with Bioanalyzer (Agilent). Samples of extracted RNA were sequenced with the Smart-Seq2 protocol for the low RNA abundance, providing datasets of unstranded, paired-end (2×101) reads of sizes between 54 Mio and 102 Mio. Smart-Seq2 adapters and poly-A tails were aggressively cut from both ends of the paired-end reads with Cutadapt v1.16, discarding reads shorter than 15 nucleotides left datasets of sizes between 37 Mio and 80 Mio. Reads were mapped to the human genome version GRCh38.p13 with splice-aware mapper STAR v2.7.3. with high alignment rates (93.3% to 94.9%) and the number of uniquely mapping, unstranded fragments per gene were determined with STAR option – quantMode GeneCounts.

These fragment counts per gene (sum over all genes 28 Mio to 63 Mio) were analyzed for differential gene expression with R v3.5.1 and DESeq2 v1.22.1 (Love et al., 2014). Normalizing with DESeq2 sizeFactors for the different dataset sizes and applying PCA to the regularized-log-transformed counts, we identify one outlier sample gfpp_g1_2 whose location in the PC1-PC2 plot is far from its biological replicates. Thus, we discarded this outlier sample from any further analysis. To be considered as differentially expressed genes comparing any two groups of samples, we required that these genes had at least 10 counts in at least one sample, an absolute log2-fold change larger than 1, and an adjusted p-value of max 0.05. Gene set enrichment analyses were performed with fgsea v1.8.0 and MSigDB v7.2.1 using an adjusted p-value cutoff of 0.05 following the method described for the single-cell RNA-seq data analysis (see above).

### Correlating live-imaging and sequencing data

FACS measurements of GFP in sequenced cells were used to link between transcriptome and live-cell data statistically. Due to potential differences between FACS and live imaging in terms of sensitivity and dynamic range, the raw GFP measurements of both methods are not directly comparable. To account for this, GFP measurements obtained from live imaging were quantile-transformed to match the distribution of GFP values corrected for forward scatter (FSC) obtained from the entire distribution of cells through flow cytometry. This strategy assumes that the ordering of GFP values revealed by both techniques is approximately equal. The GFP values of sequenced cells obtained from FACS were then grouped according to the three transcriptional clusters identified using the Leiden algorithm. For each cluster *c*, we estimated the corresponding conditional distribution *p* (*y* | *c*) over GFP levels *y* using Gaussian mixture modeling. Each cell *i* tracked via imaging was then classified into the different clusters by picking the cluster for which *p* (*y_i_* | *c*) for all *c* was maximal. Different features revealed from imaging were then analyzed according to the predicted clusters. Analyses were performed in Matlab using a custom code.

### Quantification and statistical analysis

Statistical significance tests were performed using the stat_compare_means function in the ggpubr R package v0.4.0 (https://github.com/kassambara/ggpubr). Significance was defined as *P < 0.05, **P < 0.01, ***P < 0.001, ****P < 0.0001. P values were determined by Wilcoxon and Kruskal-Wallis tests. The number of replicates is provided in the figure legends. *N* denotes the number of independent experiments, and *n* denotes the total number of measurements.

## Data availability

The raw single-cell and bulk RNA sequencing datasets have been deposited to ArrayExpress (ArrayExpress < EMBL-EBI) with the identification numbers E-MTAB-11208 and E-MTAB-11210 respectively.

