## Supplementary figures for "A combined transcriptional and dynamic roadmap of single human pancreatic endocrine progenitors reveals proliferative capacity and differentiation continuum"

### Title:

### Supplementary information:

##### Supplementary Figure 1 (Related to Figure 1):

**Further validation of the dual hESC-based reporter line monitoring human *NEUROG3* transcription and protein**
**S1A:** Representative karyograms of the dual *NEUROG3* reporter line at passage numbers 14 and 17, showing no karyotypic abnormality.
**S1B:** Representative plots showing flow cytometry gating strategy used to identify GFP+ cells. After all cells at Stage 4 Day 1 were gated by their size (first panel), singlets (second panel) were selected. Out of the singlets, live cells without live/dead stain (third panel) were analyzed for GFP signal (fourth panel).
**S1C:** Boxplots showing the comparison of the differentiation efficiency of the dual *NEUROG3* reporter line with the mother H1 hESC line, analyzed at Stage 5 Day 3 of the *in vitro* pancreatic differentiation by flow cytometry. Graph showing percentage of positive

cells over all cells stained for C-peptide and glucagon expression. *N* = 3 and n = 11 for the dual *NEUROG3* reporter line, and *N* = 1 and n = 3 for the H1 hESC line.

#### Supplementary Figure 2 (Related to Figure 2):

**Live imaging of *NEUROG3*+ human pancreatic cells reveals heterogeneity of *NEUROG3* expression and peak**
**S2A:** GFP and RFP fluorescence tracing plots for all cells that have been traced. Fluorescence intensities were scaled between 0 and 1. *N* = 3, *n* = 52
**S2B:** Raw GFP (left panels) and RFP (right panels) fluorescence traces are given separately for all cells that have been traced. Y-axes indicate the intensities in mean gray value. Numbers on top of the plots represent the traced cell ID number. *N* = 3, *n* = 52.

#### Supplementary Figure 3 (Related to Figure 3):

**Characterization of cycling human *NEUROG3*+ progenitors**
**S3A:** Location of traced cells shown over the field of view, faceted by division and colored by time.
**S3B:** Single fluorescence tracing plots of dividing and non-dividing cells, shown in light and dark colors, respectively. Plots show GFP or RFP fluorescence intensities over time for non-dividing cells or until the division event for the dividing cells. *N* = 3, *n* = 78
**S3C:** Comparison of the average movement speed of dividing and non-dividing cells. Grey dots indicate outliers of the boxplot, and black dots the actual values. P-value was determined by the Wilcoxon test.
**S3D:** Flow-cytometry-based live cell cycle analysis of cells from different stages of *in vitro* differentiation protocol. Bars indicate mean percentage values +- SEM. *N* = 2 for S3D2 and *N* = 4 for all the other groups.
**S3E:** Representative immunofluorescence microscopy images showing dividing NEUROG3+ cells at S4D1 and S4D3 of the *in vitro* pancreatic differentiation protocol (arrowheads). Anti-NEUROG3 (cyan), anti-RFP (magenta), anti-GFP (green), and anti-KI67 (gray). Scale bars, 10 µm.
S: Stage, D: Day

#### Supplementary Figure 4 (Related to Figure 4):

**Deep single-cell sequencing of *NEUROG3*+ cells indicates a gradual differentiation trajectory prior to hormonal transcription**
**S4A:** Table showing the number of *NEUROG3*+ and *NEUROG3*- cells per FACS-based GFP category.
**S4B:** Normalized gene expression levels of selected genes projected onto the UMAP.
**S4C:** Normalized gene expression levels for top 10 marker genes for each identified cluster projected onto the UMAP.
**S4D:** Table showing sequencing-based cell cycle category per sequencing clusters for a total number of 380 cells.
**S4E:** Normalized gene expression levels of selected cell cycle exit-related genes projected onto the UMAP.

#### Supplementary Figure 5 (Related to Figure 5):

**Correlating live imaging and single-cell sequencing of early hESC-derived endocrine progenitors**

**S5A:** GFP fluorescence intensities of sequenced GFP+ cells (black) and tracked cells (orange) according to their predicted or measured EP clusters.

**S5B:** Mean GFP and RFP intensities of traced cells over time, grouped by their predicted transcriptomic clusters.

**S5C:** Location of tracked non-dividing cells shown over the field of view, faceted by their predicted transcriptomic clusters. The upper panel shows tracks colored by time, and the lower panel shows tracks colored by speed (µm/hour).

#### Supplementary Figure 6 (Related to Figure 6):

**Heterogeneity of *NEUROG3* expression and peak in the live imaging of *NEUROG3*+ human pancreatic cells are also present in the 3D culture system**

**S6A:** GFP and RFP fluorescence tracing plots for all cells that have been traced from the 3D culture. Fluorescence intensities were scaled between 0 and 1. $N = 1$, $n = 30$.

**S6B:** Raw GFP and RFP fluorescence traces are given separately for all cells traced from the 3D culture. Y-axes indicate the intensities in mean gray value. $N = 1$, $n = 30$ Numbers on top of the plots represent the traced cell number.

#### Supplementary Figure 7 (Related to Figure 7):

**Dividing *NEUROG3*+ cells are in transition to the endocrine progenitor lineage**

**S7A:** GSEA of GFP+ G2M cells compared with GFP+ G0/G1 cells.

Figure S1

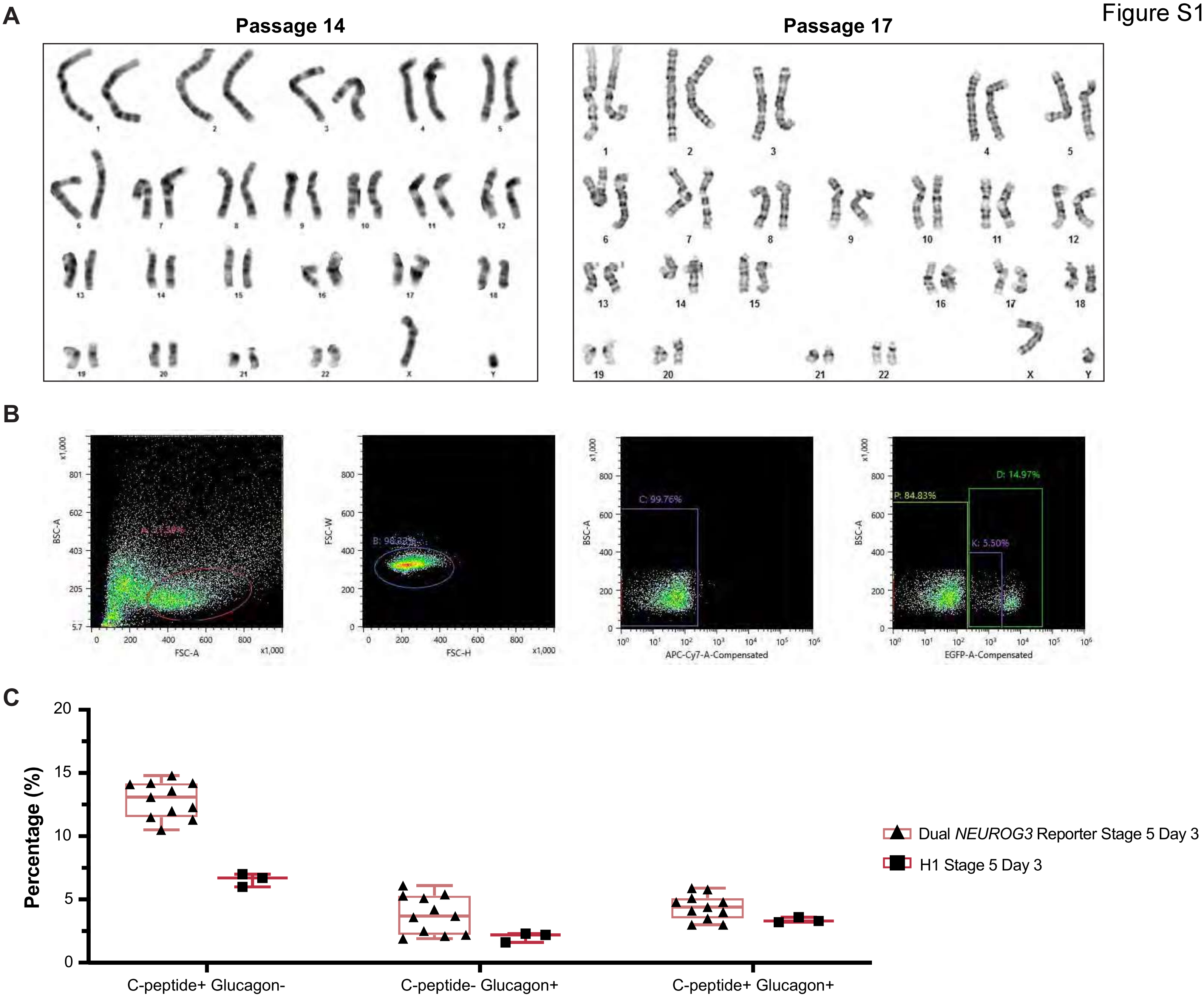

A
Passage 14
Passage 17
B
BSC-A
FSC-A
FSC-W
FSC-H
APC-Cy7-A-Compensated
EGFP-A-Compensated
C: 99.76%
P: 84.83%
D: 14.97%
K: 5.50%
C
Percentage (%)
C-peptide+ Glucagon-
C-peptide- Glucagon+
C-peptide+ Glucagon+
Dual *NEUROG3* Reporter Stage 5 Day 3
H1 Stage 5 Day 3

Figure S2

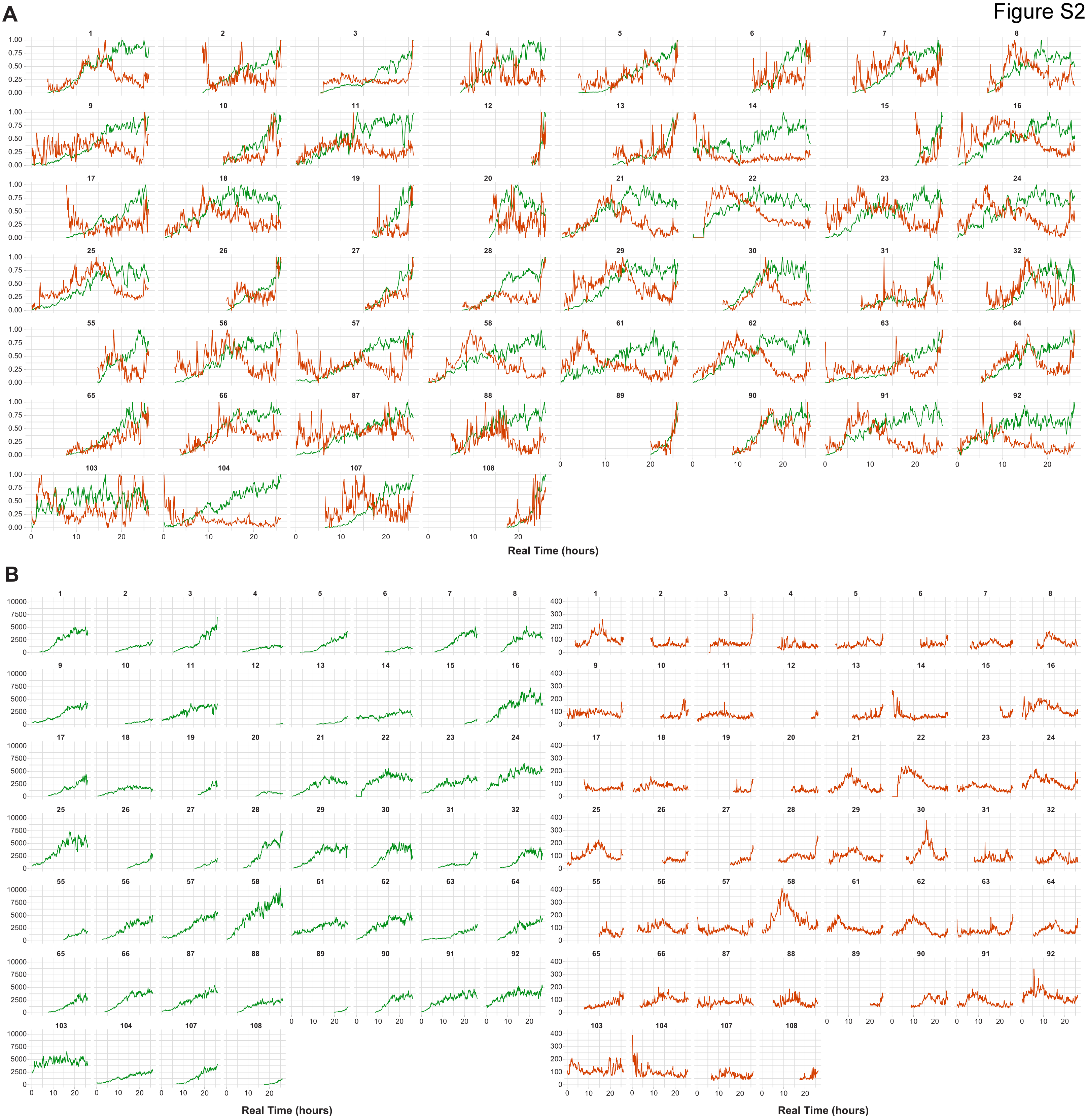

Figure S3

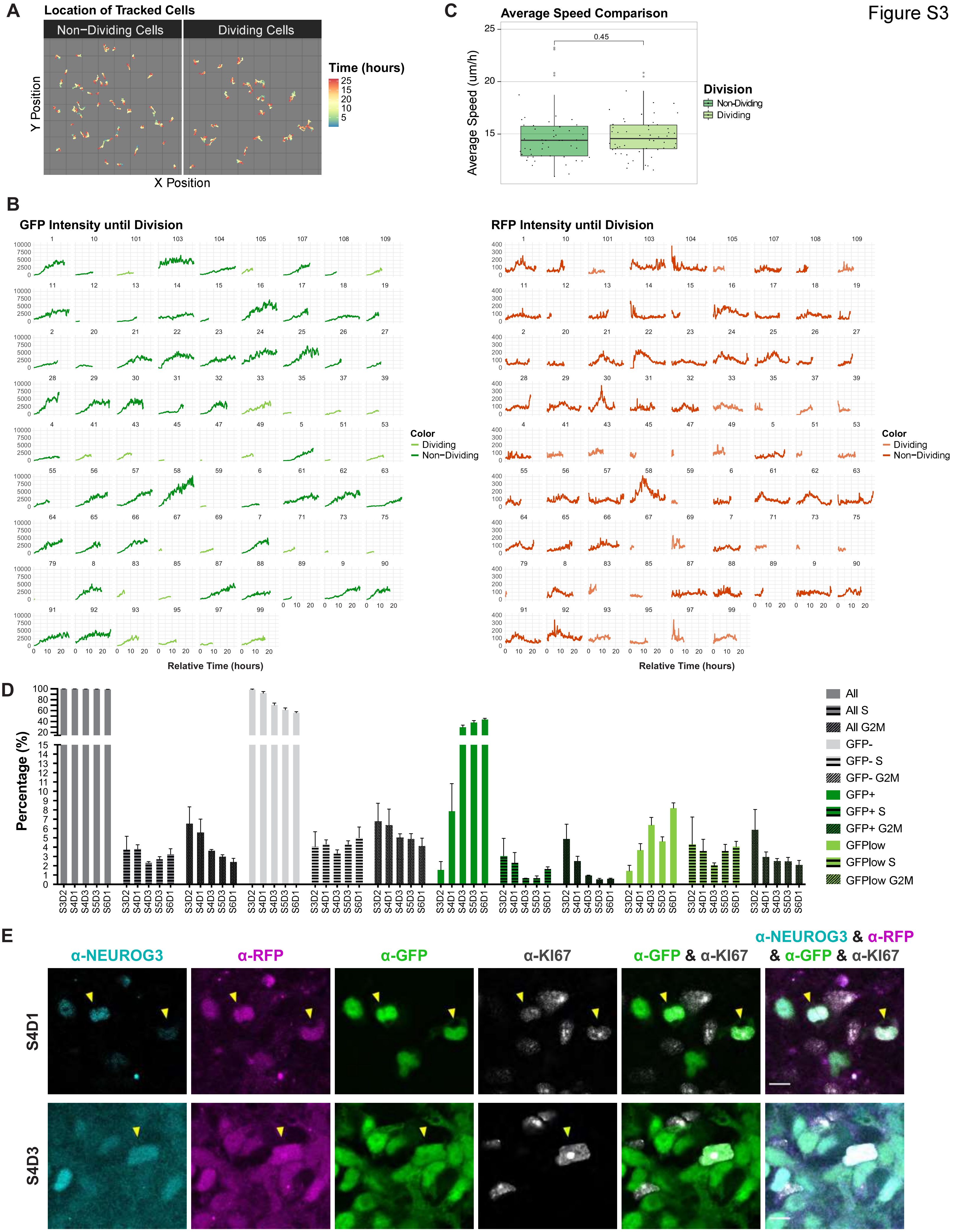
A
Location of Tracked Cells
Non−Dividing Cells
Dividing Cells
Y Position
X Position
Time (hours)
25
15
20
10
5
C
Average Speed Comparison
0.45
Average Speed (um/h)
25
20
15
Division
Non-Dividing
Dividing
B
GFP Intensity until Division
RFP Intensity until Division
Color
Dividing
Non−Dividing
Relative Time (hours)
D
Percentage (%)
All
All S
All G2M
GFP-
GFP- S
GFP- G2M
GFP+
GFP+ S
GFP+ G2M
GFPlow
GFPlow S
GFPlow G2M
S3D2
S4D1
S4D3
S5D3
S6D1
E
α-NEUROG3
α-RFP
α-GFP
α-KI67
α-GFP & α-KI67
α-NEUROG3 & α-RFP & α-GFP & α-KI67
S4D1
S4D3

Figure S4

**A**

| | GFP- | GFP low | GFP high | All |
|---|---|---|---|---|
| ***NEUROG3-*** | 85 | 58 | 44 | 187 |
| ***NEUROG3+*** | 6 | 136 | 51 | 193 |

**B**

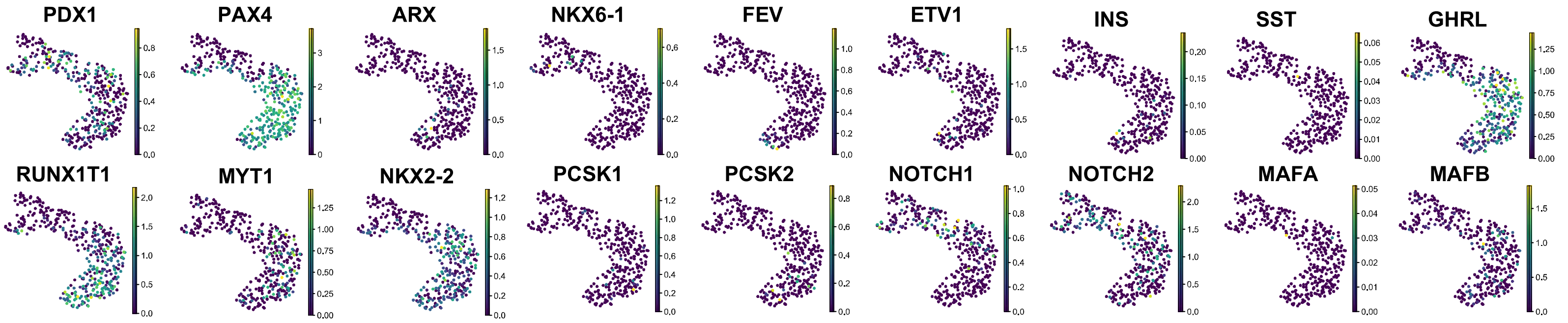

**C**

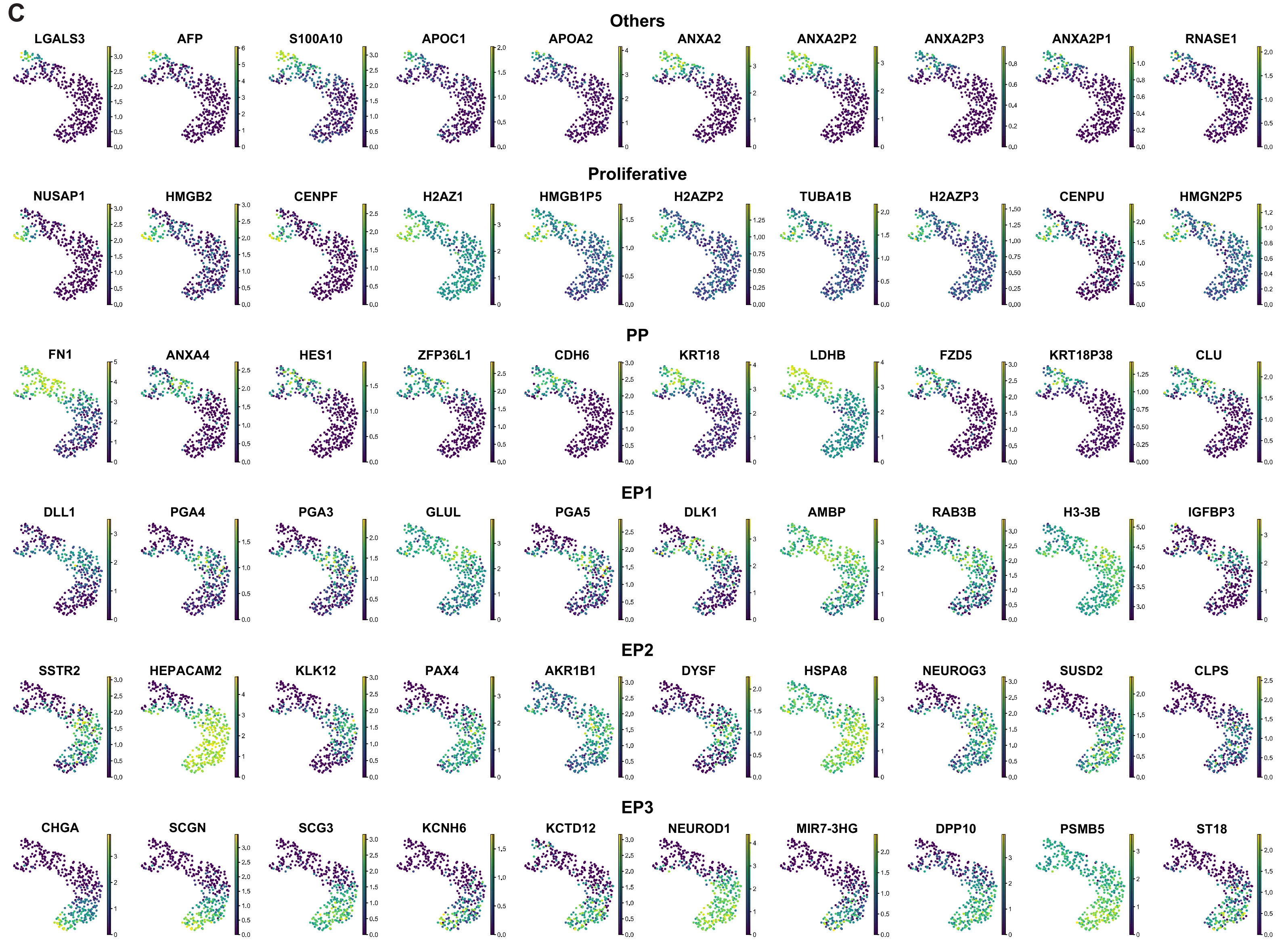

**D**

| Cluster | G1 | S | G2M |
|---|---|---|---|
| Others | 64,0% | 24,0% | 12,0% |
| Proliferative | 0,0% | 21,9% | 78,1% |
| PP | 59,0% | 23,0% | 18,0% |
| EP1 | 81,3% | 7,8% | 10,9% |
| EP2 | 77,9% | 7,4% | 14,7% |
| EP3 | 93,2% | 3,9% | 2,9% |

**E**

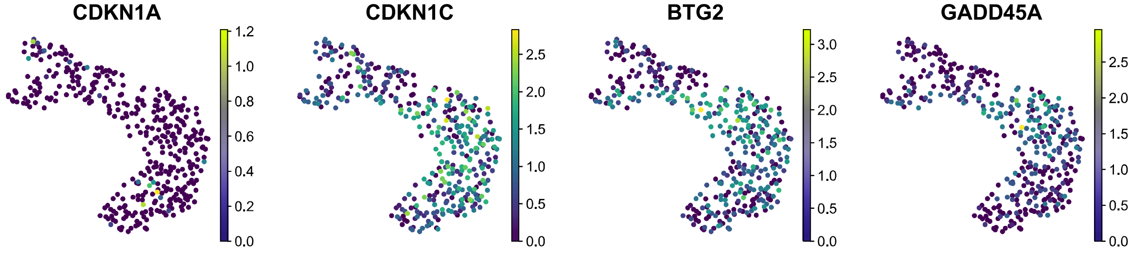

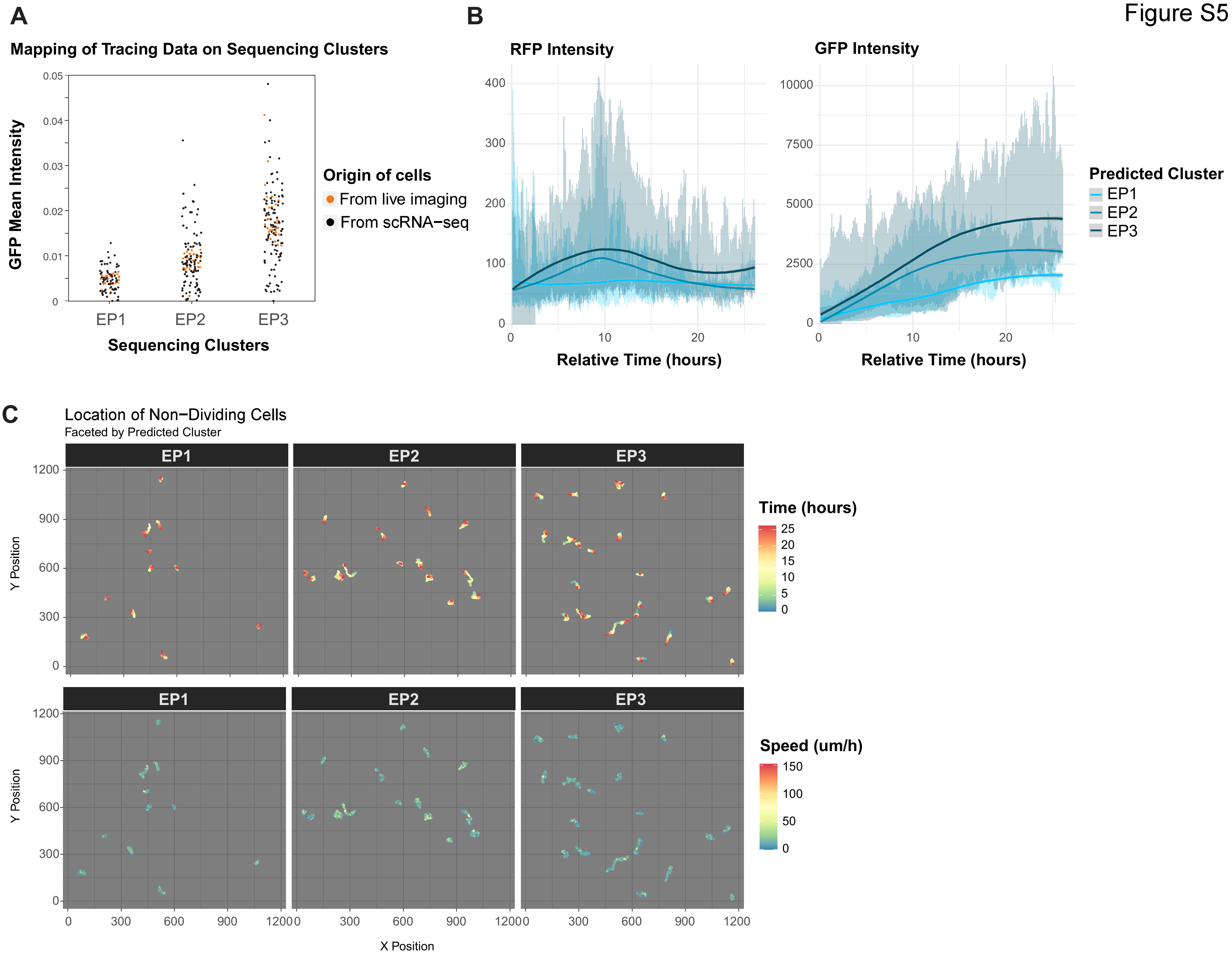
A
Mapping of Tracing Data on Sequencing Clusters
GFP Mean Intensity
Sequencing Clusters
EP1
EP2
EP3
Origin of cells
From live imaging
From scRNA−seq
B
RFP Intensity
GFP Intensity
Relative Time (hours)
Predicted Cluster
EP1
EP2
EP3
C
Location of Non−Dividing Cells
Faceted by Predicted Cluster
Y Position
X Position
Time (hours)
Speed (um/h)

Figure S6

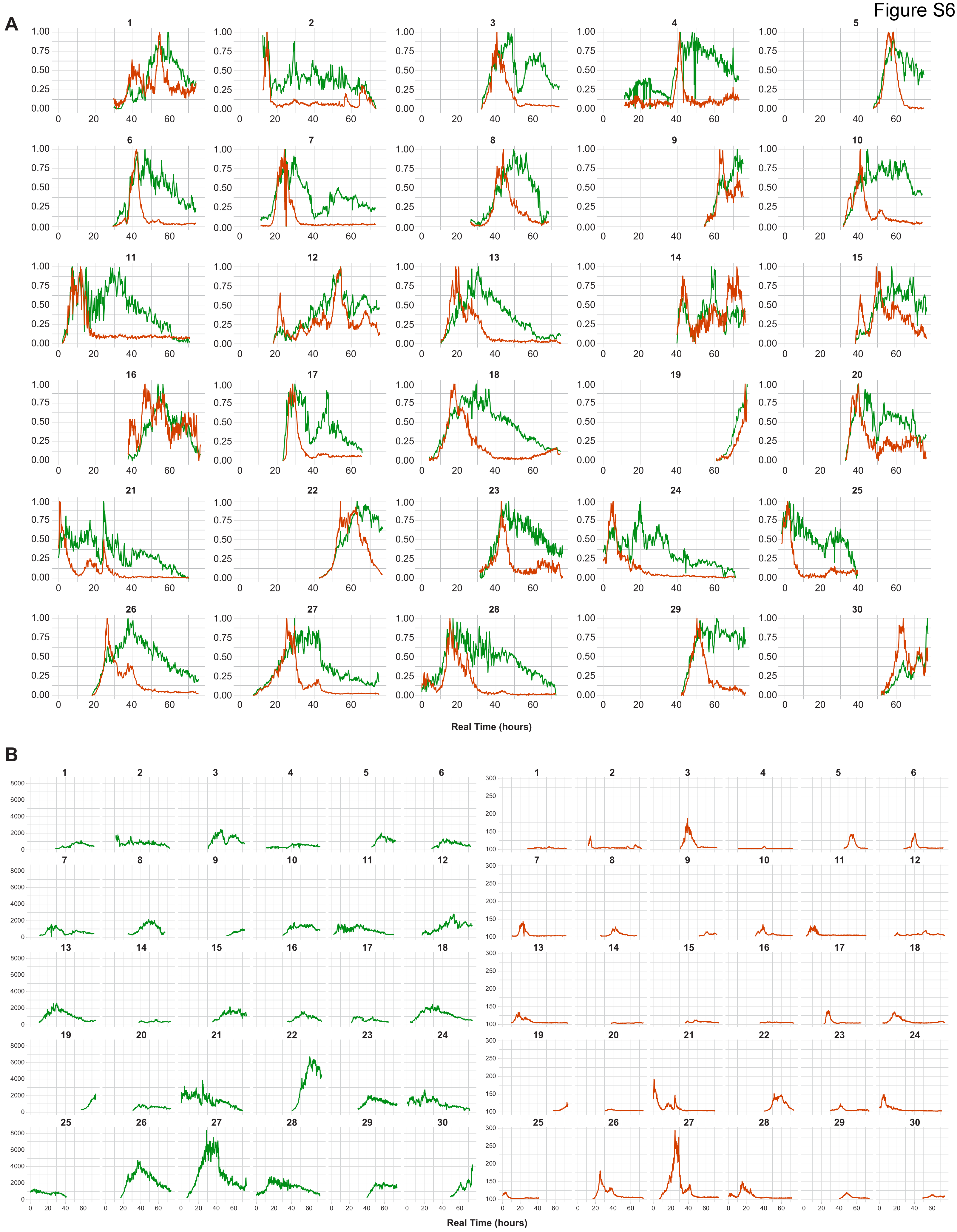
A
0.00
0.25
0.50
0.75
1.00
0
20
40
60
Real Time (hours)
B
0
2000
4000
6000
8000
100
150
200
250
300
Real Time (hours)

Figure S7

**A**

GFP+ G2M — GFP+ G0/G1

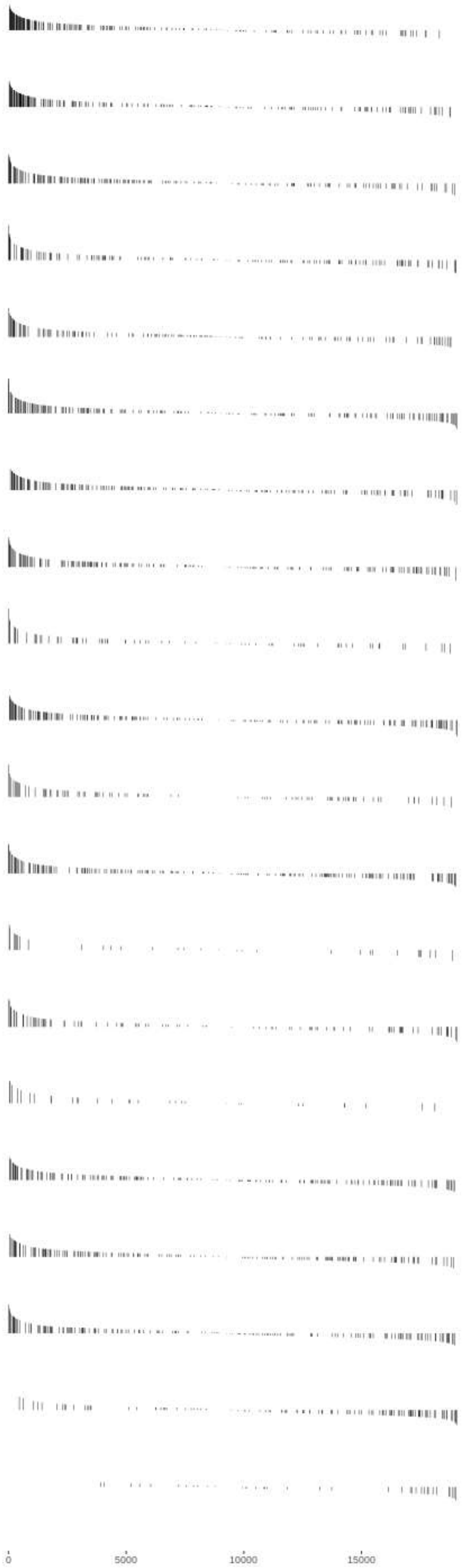

| Pathway | Gene Ranks | NES | p Value | p Adjusted |
|---|---|---|---|---|
| HALLMARK_E2F_TARGETS | | 2.08 | 1.0e-04 | 1.7e-03 |
| HALLMARK_G2M_CHECKPOINT | | 1.98 | 1.0e-04 | 1.7e-03 |
| HALLMARK_TNFA_SIGNALING_VIA_NFKB | | 1.75 | 1.0e-04 | 1.7e-03 |
| HALLMARK_APOPTOSIS | | 1.68 | 2.1e-04 | 2.2e-03 |
| HALLMARK_UV_RESPONSE_DN | | 1.70 | 2.2e-04 | 2.2e-03 |
| HALLMARK_EPITHELIAL_MESENCHYMAL_TRANSITION | | 1.62 | 3.1e-04 | 2.6e-03 |
| HALLMARK_MITOTIC_SPINDLE | | 1.58 | 4.1e-04 | 2.6e-03 |
| HALLMARK_IL2_STAT5_SIGNALING | | 1.58 | 5.2e-04 | 2.9e-03 |
| HALLMARK_CHOLESTEROL_HOMEOSTASIS | | 1.67 | 2.2e-03 | 1.0e-02 |
| HALLMARK_HYPOXIA | | 1.50 | 2.6e-03 | 1.0e-02 |
| HALLMARK_INTERFERON_ALPHA_RESPONSE | | 1.63 | 2.7e-03 | 1.0e-02 |
| HALLMARK_P53_PATHWAY | | 1.49 | 2.9e-03 | 1.0e-02 |
| HALLMARK_NOTCH_SIGNALING | | 1.70 | 3.7e-03 | 1.2e-02 |
| HALLMARK_ANDROGEN_RESPONSE | | 1.57 | 4.4e-03 | 1.3e-02 |
| HALLMARK_ANGIOGENESIS | | 1.69 | 4.5e-03 | 1.3e-02 |
| HALLMARK_KRAS_SIGNALING_UP | | 1.47 | 5.4e-03 | 1.5e-02 |
| HALLMARK_INTERFERON_GAMMA_RESPONSE | | 1.47 | 5.6e-03 | 1.5e-02 |
| HALLMARK_INFLAMMATORY_RESPONSE | | 1.47 | 6.1e-03 | 1.5e-02 |
| HALLMARK_UNFOLDED_PROTEIN_RESPONSE | | -1.79 | 1.1e-03 | 5.6e-03 |
| HALLMARK_PANCREAS_BETA_CELLS | | -2.23 | 4.1e-04 | 2.6e-03 |
